# EPHB2 promotes long-range axon extension and its variants cause callosal dysgenesis

**DOI:** 10.64898/2026.11.13.766231

**Authors:** Sung Soon Park, Gergely N. Nagy, Isabel Pérez Ferrer, Yaiza Coca, Macarena Herrera, Camille Kahwaji, Xiaolei Brian Zhang, Lang Liu, Srikanth Kolluru, Mauricio Neira, Yacine Kerdjemil, Flavia Palombo, Chiara La Morgia, Leonardo Caporali, Valerio Carelli, Quinten Waisfisz, Mariet W. Elting, Gunnar Douzgos Houge, Gyri Aasland Gradek, Dag Tveitnes, Eamonn G. Sheridan, Katrina Prescott, Melody Redman, Aliza Imam, Pia Ostergaard, Katelynn Anderson, David D. Weaver, Elise Schaefer, Vincent Laugel, Ziv Gan-Or, Matthew B. Dalva, Christel Depienne, E. Yvonne Jones, Eloisa Herrera, Artur Kania

## Abstract

The corpus callosum (CC) is the largest fibre tract connecting the cerebral hemispheres, essential for integrating neural activity across large-scale brain networks. Accordingly, abnormal CC development is associated with diverse neurodevelopmental and neuropsychiatric disorders. Callosal projection neurons (CPNs) of the cortex form the CC by extending long-range axons to the contralateral hemisphere; however, the mechanisms that sustain axon elongation over such remarkable distances remain poorly understood. Here we show that variants in the human gene *EPHB2*, which encodes a tyrosine kinase receptor regulating nervous system development and implicated in neurodevelopmental disorders, cause CC dysgenesis. We characterise variant-specific effects on EPHB2 signalling and demonstrate that variants identified in individuals with distinct CC morphologies differentially alter CPN axon outgrowth and integration in vivo. We further show that EphB2 has a cell-autonomous, ligand-independent function in promoting CPN axon outgrowth by spatially organising actomyosin at axonal growth cones. These findings reveal a previously unrecognised role for EphB2 in promoting long-range axon extension and provide the first evidence that *EPHB2* variants cause CC dysgenesis in humans. More broadly, our study links molecular variation in *EPHB2* to individualised abnormalities in brain connectivity and provides a framework for understanding how heterogeneous circuit-level phenotypes arise in neurodevelopmental disorders.

## Introduction

A defining feature of eutherian mammalian brains is the corpus callosum (CC), the largest commissural fibre tract connecting the cerebral hemispheres^1^. By coordinating the exchange and integration of neuronal activity between the hemispheres, the CC contributes to large-scale brain network function underlying higher-order cognitive processes including learning, language processing, social cognition, and multisensory integration^2–4^. Accordingly, callosal dysgenesis, characterised by partial or complete absence, or abnormal morphology of the CC, is associated with a broad spectrum of neuropsychiatric and neurodevelopmental disorders^5–8^.

One feature that distinguishes callosal projection neurons (CPNs), which give rise to the CC, from most other central nervous system neurons is their ability to extend extraordinarily long axons^9^. CPN development involves many processes including postmitotic cell-fate specification, acquisition of axon-dendrite polarity during somatic migration, long-distance axonal extension, crossing of the interhemispheric midline, and the establishment of synaptic connections within the contralateral cortex^10,11^. While numerous genes and associated mechanisms have been implicated in each of these developmental processes^5^, the mechanisms enabling sustained long-range growth of CPN axons remain incompletely understood.

Rare coding variants in *EPHB2* have recently been identified in individuals with neurodevelopmental syndromes^12–21^, while their pathogenic contributions are less understood^22,23^. EphB2 belongs to the Eph receptor tyrosine kinase family whose members employ diverse signalling modes to regulate development in multiple tissues^24^. Canonical Eph signalling is initiated by binding with its ligand ephrin on the opposing cell surface, followed by receptor clustering, tyrosine kinase activation, and ephrin-Eph receptor trans-endocytosis. In neurons, Eph signalling primarily induces cytoskeletal collapse and growth cone repulsion, contributing to neuronal soma migration and neurite targeting during neural circuit formation^25^. Genetic studies in mice have established an essential role for EphB2 in CC development^26–29^. However, the primary cellular function of EphB2 during CPN development remains unclear, partly owing to its widespread and dynamic expression in the developing nervous system^26,30^.

Neuronal growth cones undergo context-dependent conformational changes that reflect distinct modes of axon behaviour during development^31^. Growth cone morphology and motility are shaped by the dynamic interplay between actin cytoskeleton and associated molecular motors, including non-muscle myosin II (NMII)^32,33^. NMII activity and organisation, regulated by the RhoA-ROCK signalling pathway within growth cones, are crucial for long-range axon extension^34–37^. However, the upstream mechanisms that spatially organise these effectors within growth cones remain largely unknown.

Here, we investigated the molecular and cellular consequences of human *EPHB2* variants identified in individuals with CC dysgenesis and other neurodevelopmental defects. By integrating clinical and genetic analyses with cell-based functional assays, structural modelling, and CPN-specific *in vivo* and *in vitro* approaches, we examined how variant-specific effects on EPHB2 function contribute to CC development. We identify a dosage-sensitive, cell-autonomous requirement for EphB2 during CPN axon elongation. We further show that EphB2 promotes axon outgrowth through a ligand-independent signalling mode involving spatial organisation of actomyosin through interaction with RhoA guanine nucleotide exchange factors (GEFs) within growth cones. These findings establish EphB2 as a key regulator of CC morphology and uncover an unexpected signalling mode through which Eph receptors can promote long-range axon extension.

## Results

### Individuals carrying rare *EPHB2* coding variants display corpus callosum dysgenesis and other neurodevelopmental defects

A cohort of eight probands harbouring variants within the coding region of the *EPHB2* gene was identified through whole exome sequencing (WES) of individuals exhibiting early onset neuroanatomical and neurological defects (Fig.1a,b). The cohort includes three individuals with *de novo* heterozygous variants, two individuals with inherited compound heterozygous variants and three siblings sharing the same set of compound heterozygous variants. All variants are absent or extremely rare (gnomAD v.4.1.1), and most are predicted by multiple *in silico* tools to have deleterious effects on EPHB2 functions (e.g. all have combined annotation dependent depletion score > 20) (Extended Data Fig. 1a). Except for the frameshift variant identified in proband 1, which is predicted to introduce a premature stop codon early in the coding sequence, all other eight variants are missense. The *EPHB2* gene is extremely constrained and highly intolerant of genetic variants (missense Z-score = 4.24; pLI = 1). All affected residues, except for the frameshift variant p.V268Sfs*64 (p.V268Sfs hereafter), are in sequences highly conserved among *EPHB2* homologues from other species and Eph receptor family members (Fig. 1c,d). The variants are located in all known EPHB2 protein domains, which could cause diversified molecular impacts and phenotypic presentation.

**Fig. 1:**
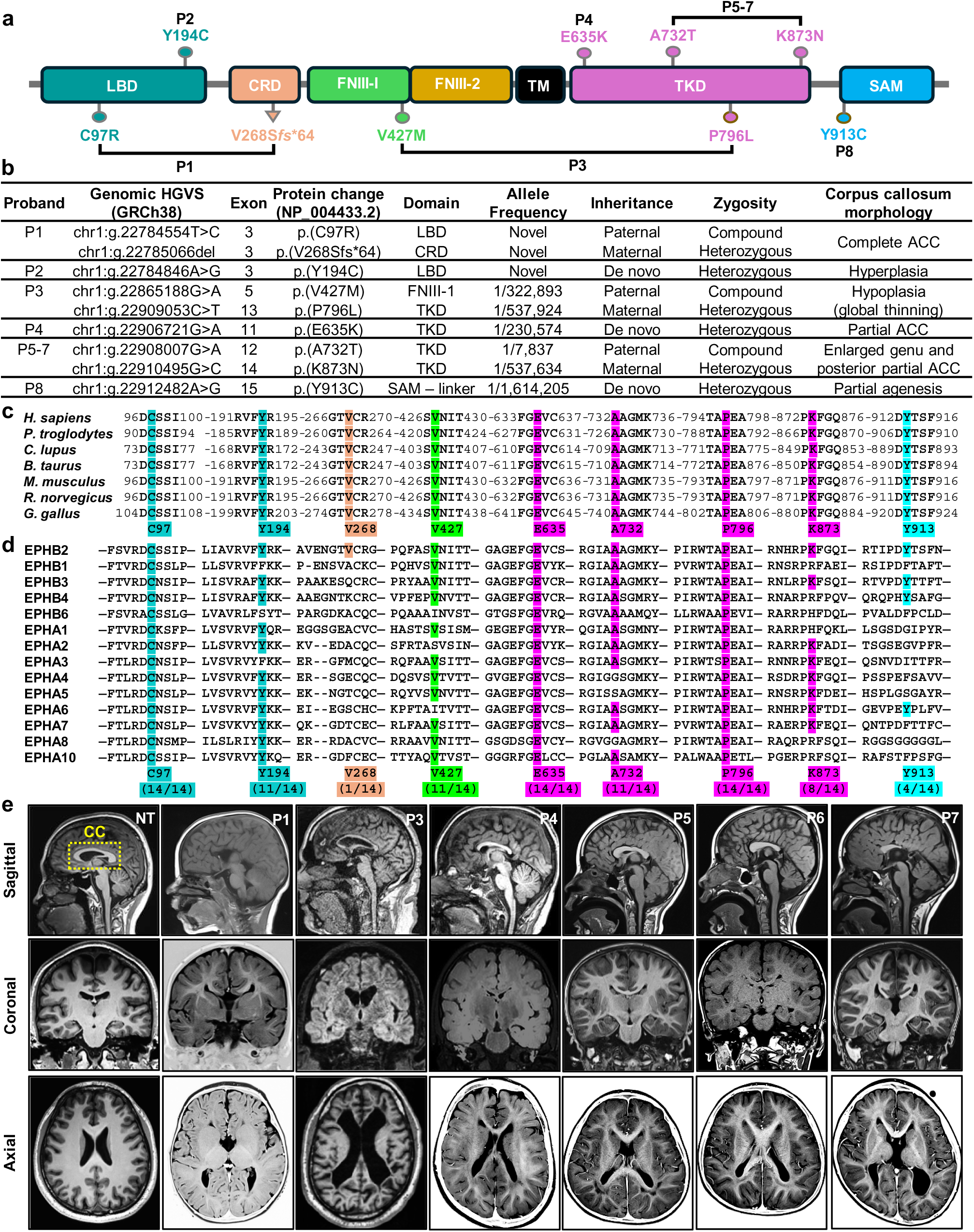
Callosal dysgenesis in probands carrying *EPHB2* variants. **a**, Location of human *EPHB2* variants within the protein coding sequence and domains (colour coded) of EPHB2 in probands P1-8. Probands trans-heterozygous for two variants are shown as horizontal lines. Variants are colour coded according to their EPHB2 domain location. LBD: ligand-binding domain, CRD: cysteine-rich domain (includes sushi domain and epidermal growth factor-like (EGF-like) domain), FNIII-1/2: fibronectin type III domain 1 and 2, TM: transmembrane region, TKD: tyrosine kinase domain; SAM, sterile-alpha motif domain. Human EPHB2 isoform 2 (NP_004433.2) is shown. **b,** Table summarising the molecular and clinical profile of EPHB2 variant probands. Genomic coordinates and variants were annotated based on the human reference genome GRCh38 (hg38, NCBI Accession NC_000001.11 for chromosome 1). Exon number is based on EPHB2 transcript isoform 2 (NM_004442.7). **c,d,** EPHB2 protein sequences spanning the variant residues are conserved between orthologous (b) and between human EPH paralogues (c). Variant residue numbers at the bottom of the alignment correspond to the human EPHB2 sequence. The number in parenthesis indicates the conservation rate across human paralogues. **e,** Brain magnetic resonance imaging (MRI) scans of a neurotypical individual (NT) and probands carrying *EPHB2* variants where images were available in the clinical report. The corpus callosum (CC) is bounded by the yellow dashed box.

The cohort originates from six different families with diverse ancestral background across North America and Europe with all probands exhibiting a spectrum of neurodevelopmental and neuropsychiatric manifestations including global developmental delay and intellectual disability, aphasia, repetitive behaviour, anxiety, and attention–deficit/hyperactivity disorder (ADHD) (Extended Data Fig. 1b). Visual impairment was present in four individuals and ranged from mildly reduced visual acuity to central blindness. One individual present deafness. Additional nervous system findings include seizures, and some individuals had fine motor developmental delay, hypotonia, or poor coordination. Dysmorphic features were variable but recurrent, including narrow forehead, abnormal ears, and broad nasal bridge, characteristic of neurodevelopmental disorders.

Importantly, all individuals within the cohort show CC dysgenesis of varying forms and severities based on brain magnetic resonance imaging (MRI) reports (Fig. 1b). Proband 1 exhibits a complete CC agenesis, proband 2 has an enlarged splenium, proband 3 shows a global CC thinning, proband 4 shows partial agenesis of the rostrum and splenium, proband 5 through 7, who are siblings with the same heterozygous variants, show partial agenesis of the splenium and rostrum and with a shortened callosal body and blunted genu, and proband 8 shows partial agenesis in the splenium (Fig. 1e; images unavailable for probands 2 and 8). Additional neuroimaging findings include cerebellar atrophy (probands 5-7), microcephaly (probands 2, 4, 5-7) and abnormalities of cortical and white matter development (probands 3, 4, 5-7).

We next asked whether *EPHB2* coding sequence variants are also associated with morphological heterogeneity of CC in populations beyond our cohort. To address this, we conducted a rare variant genetic association analysis between *EPHB2* variants and the average volume of CC in individuals in the UK Biobank (UKB)-derived cohort for which T1-weighted brain MRI data are available. Over 60,000 individuals were included in the final analysis after excluding individuals with neurological conditions that may have confounding effects on CC volume (refer to methods for exclusion criteria.). The cohort was divided into two groups: 53,919 neurotypical individuals and 7,617 individuals with reported mental health phenotypes, following the observation from our own cohort of patients who all present with comorbid mental disorders (see Extended Data Fig. 1b). Interestingly, a gene-based rare variant analysis identified a significant association between rare nonsynonymous *EPHB2* variants, but not noncoding variants, and average CC volume in individuals with mental health phenotypes (SKAT-O p = 0.034; Extended Data Fig. 1c). No association was observed using a burden test (p = 1.0), suggesting that the observed signal is driven by heterogeneous variant-specific effects rather than a uniform directional effect across variants. No evidence of association was observed in neurotypical controls (SKAT-O p = 0.528; burden p = 1.0). Collectively, these findings suggest that rare coding variation in *EPHB2* (full list of UKB cohort variants available in Supplementary Table 1) may contribute to variation in CC morphology in individuals with neurodevelopmental or neuropsychiatric conditions.

### *EPHB2* variants impair specific steps in EPH signalling

Next, we asked whether the identified human variants have deleterious impacts on EPHB2 protein functions. To investigate this, we assessed the impact of all 9 variants on each step of the Eph receptor canonical signalling cascade (Fig. 2a). First, we evaluated ligand-binding by expressing EPHB2 variants as GFP-fused proteins in HeLa cells that lack endogenous B-type EPH receptors, exposed them to pre-clustered ephrin-B1 (eB1) conjugated with Cy5 for 30 minutes and quantified eB1 cellular accumulation. Cells expressing wild type EPHB2 (WT-EPHB2) showed a robust accumulation of eB1 relative to the receptor expression level measured by GFP intensity (Fig. 2b). In contrast, expression of EPHB2 lacking ligand-binding domain (ΔLBD-EPHB2), as an interaction-abolished control, did not show any ligand retention regardless of receptor expression level (Fig. 2c). Interestingly, the C97R variant significantly impaired eB1 accumulation, while V268Sfs failed to localise to the cell surface and was unable to accumulate eB1. Other EPHB2 variants displayed ephrin retention comparable to WT-EPHB2. To further confirm this observation, we performed biolayer interferometry (BLI) by titration of immobilised EPHB2 ectodomain with the receptor binding domains (RBD) of ephrin-B1 and ephrin-B2. While both ligands exhibited high-affinity binding to the WT-EPHB2 ectodomain, neither ephrin-B RBD showed detectable binding to C97R-EPHB2 or ΔLBD-EPHB2 (Extended Data Fig. 2a). Together, these results indicate that the C97R and V268Sfs variants significantly impair ephrin-B ligand binding.

**Fig. 2:**
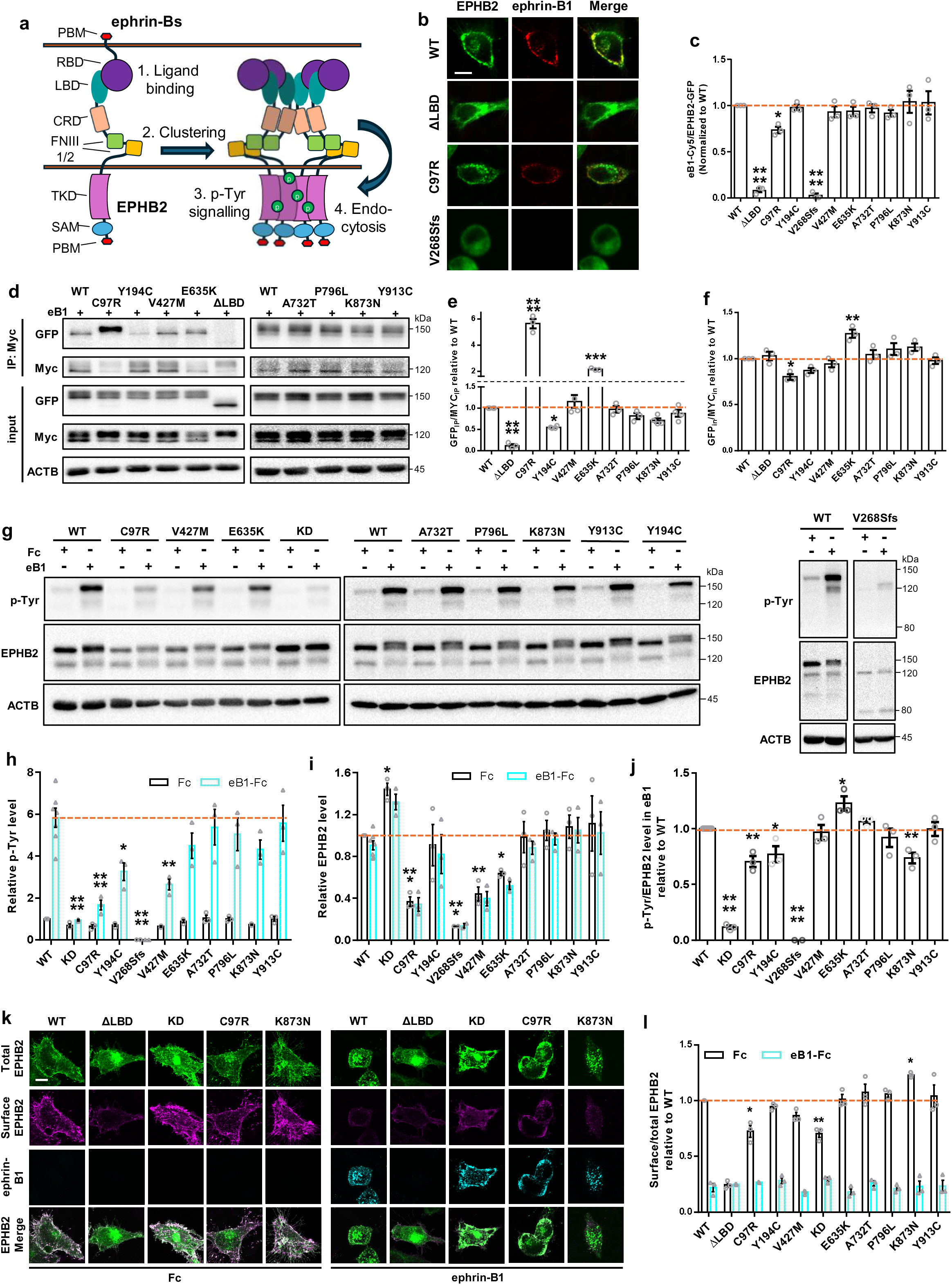
EPHB2 variants impair distinct EPHB2 signalling steps. **a**, Schematic diagram of the canonical EphB2 receptor signalling cascade. PBM: postsynaptic density-95/Discs large/zona occludens-1 (PDZ)-binding motif, RBD: receptor-binding domain, LBD: ligand-binding domain, CRD: cysteine-rich domain, FNIII-1/2: fibronectin type III domain 1 and 2, TKD: tyrosine kinase domain; SAM, sterile-alpha motif domain. Ephrin ligand binding elicits receptor-ligand complex clustering, causing activation of the tyrosine kinase, resulting in substrate phosphorylation and autophosphorylation. Signalling termination involves complex endocytosis and degradation. **b,** Evaluation of variant EPHB2 ligand binding efficiency by overlay of EPHB2-expressing HeLa cells with ephrin-B1 ligand. Each row shows the same cell. Left panels: EPHB2 variants fused with GFP detected with anti-GFP antibodies. Middle panels: detection of ephrin-B1 via its conjugated Cy5 fluorophore. The right panels show merged images. Scale bar, 10 µm. **c,** Quantification of relative ephrin-B1-Cy5 mean intensity normalised to EPHB2-GFP mean intensity in cells expressing EPHB2 variants following ephrin-B1 treatment for 30 minutes. SEM; N>3 (15∼20 cells in each replicate). **d,** Immunoblots of immunoprecipitation (IP) and input lysates from the clustering assays. GFP-tagged EPHB2 variants were co-immunoprecipitated with Myc-tagged WT-EPHB2. **e,f,** Quantification of variant-GFP levels normalised to WT-Myc levels in IP (f) and input (g) lysates. SEM; N>3. **g,** Immunoblots of cell lysates from DIV6 mouse cortical neurons transfected with EPHB2 variants and treated with either Fc IgG fragment or pre-clustered ephrin-B1 for 30 minutes. **h-j,** Quantification of relative phospho-tyrosine (p-Tyr) (h), EPHB2 (i), and phospho-tyrosine/EPHB2 (j) levels in cell lysates from DIV6 mouse cortical neurons transfected with EPHB2 variants. SEM; N>4. **k,** Endocytosis assay in HeLa cells expressing wild type control, ΔLDB, C97R and K873N EPHB2 variants. Left group are cells treated with Fc IgG fragment (negative control), and the right group are cells treated with pre-clustered ephrin-B1 for 30 minutes. Image of the same cell shown in each column. Top rows show total EPHB2 detected with an anti-EPHB2 antibody, second from the top is surface EPHB2 detected with an extracellular antibody under non-permeabilised conditions, third row shows ephrin-B1 via its conjugated Cy5 fluorophore, and bottom row shows a merge of all panels above. Scale bar, 10 µm. **l,** Quantification of surface (a-ectoEphB2)/total EPHB2 (GFP) levels in HeLa cells transfected with EPHB2 variants and treated with Fc IgG fragment or pre-clustered ephrin-B1. SEM; N=3.

To assess whether the variants impact EPHB2 receptor clustering following eB1 stimulation, we co-expressed GFP-tagged EPHB2 variants with Myc-tagged WT-EPHB2 and quantified their co-immunoprecipitation (co-IP). Following eB1 treatment, WT EPHB2–GFP co-immunoprecipitated with WT EPHB2–Myc, whereas association of ΔLBD with WT EPHB2 was undetectable (Fig. 2d,e)^38–40^. Most missense variants behaved as WT-EPHB2; however, C97R and E635K showed significantly higher association while Y194C showed a significant reduction. Interestingly, we observed that E635K variant dominantly reduced the level of WT-EPHB2 in input lysates when expressed together (Fig. 2f), likely increasing their co-IP ratio. Together, we observed that C97R and Y194C alter eB1-mediated receptor clustering efficiency, whereas the effect of E635K could not be distinguished from its effect on receptor abundance.

Ephrin binding and receptor clustering allow EPHB2 receptor autophosphorylation of juxta-membrane tyrosine residues and tyrosine kinase domains further amplifying tyrosine kinase activity^41^. Therefore, we measured receptor autophosphorylation levels after transfecting GFP-tagged EPHB2 variants, distinguished from the endogenous EphB2 by their respective molecular weights of ∼150 kDa and ∼120 kDa, in mouse primary cortical neurons or HeLa cells following treatment with either Fc IgG fragment or pre-clustered eB1. In mouse primary neurons, WT-EPHB2 strongly drove autophosphorylation upon eB1 treatment compared to the Fc control (Fig. 2g,h), while a kinase-dead (KD) EPHB2 (KD-EPHB2), which lacks tyrosine kinase activity^42^, failed to respond. Interestingly, the KD-EPHB2 consistently showed higher level of basal receptor expression compared to WT-EPHB2 (Fig. 2i), consistent with the role of kinase activity in receptor turnover^43,44^. Conversely, several human variants showed reduced receptor abundance, potentially reflecting decreased protein stability or increased degradation. Interestingly, C97R and Y194C further decreased normalised receptor autophosphorylation level (p-Tyr/EPHB2) compared to WT (Fig. 2j), potentially reflecting their compromising impacts on ephrin binding and clustering, respectively. Although E635K showed lower receptor expression level, it led to an increase in normalised autophosphorylation level compared to WT, suggesting its enhancing role in tyrosine kinase activity. On the other hand, K873N did not affect receptor expression level, yet it lowered autophosphorylation levels significantly. Similar results were observed in HeLa cells (Extended Data Fig. 2b-d), together suggesting that some EPHB2 variants can perturb receptor stability and autophosphorylation.

As ligand-induced internalisation regulates EphB signalling and contact-mediated repulsion^45–47^, we examined the effects of EPHB2 variants on receptor localisation and internalisation following eB1 stimulation in HeLa cells. Under basal conditions, WT-EPHB2 receptors were strongly enriched on the cell surface (Fig. 2k left), as detected under non-permeabilising conditions using an antibody against its extracellular region. Following eB1 treatment, the bulk of WT-EPHB2 was redistributed from the cell surface to intracellular compartments, where it co-localised with eB1, accompanied by a reduction in surface receptor levels (Fig. 2k right, l), reflecting rapid ligand-induced internalisation. In contrast, most ΔLBD-EPHB2 proteins, which still preserve membrane-targeting signal peptide, were retained intracellularly under basal conditions, suggesting the key role of LBD in cell surface targeting. Upon eB1 treatment, ΔLBD-EPHB2 failed to undergo endocytosis. Interestingly, KD-EPHB2 exhibited an increase in both surface and cytosolic receptor expression compared to WT-EPHB2 (Extended Data Fig. 2e), consistent with the increase in the protein levels in cell lysates (Fig. 2g,i and Extended Data Fig. 2b,d). However, the resulting fraction of KD-EPHB2 on the cell surface was lower due to its relative enrichment in cytosol. Moreover, eB1 treatment failed to induce robust endocytosis of surface KD-EPHB2 (Extended Data Fig. 2f), consistent with the involvement of kinase activity in endocytosis^47^. Interestingly, C97R also showed a significantly reduced basal surface expression compared to WT as well as impaired endocytosis, reflecting compromised ephrin binding and subsequent signalling. In contrast, K873N showed an increase in basal surface expression without affecting endocytosis, suggesting that the reduced kinase activity may disrupt basal receptor turnover.

Ephrin-B–EPHB2 signalling was shown to cooperate with NMDA receptor (NMDAR) function to regulate callosal projection neuron (CPN) synapse development^48^. We therefore examined whether the EPHB2 variants disrupt interaction with GluN1, the obligatory NMDAR subunit that directly binds to FNIII-2 domain of EPHB2 depending on extracellular phosphorylation^42,49^. We expressed EPHB2 variants together with GluN1 in HEK293T cells and assessed their ability to form a complex via co-IP. Intriguingly, most variants, including the KD-EPHB2, exhibited comparable binding to GluN1, except for V427M (Extended Data Fig. 2g,h). Consistent with this, the Y504 extracellular phosphorylation was compromised only in the V427M variant (Extended Data Fig. 2i), suggesting that the variant may disrupt EPHB2–NMDAR interaction through inhibition of extracellular phosphorylation. Together, these biochemical and cellular findings provide functional evidence for the deleterious effects for some of the identified variants, demonstrating that they selectively disrupt specific steps of the EPHB2 receptor signalling cascade.

### Structural basis of variant EPHB2 functional defects

To provide structural context for these EPHB2 functional defects, we examined the positions and local structural environments of the variants in EPHB2 structures and models. As no experimental structure of full-length EPHB2 exists, we used EPHB2 domain structures, homologous Eph receptor structures and AlphaFold models (Fig. 3a–l, Extended Data Fig. 3). To place the extracellular variants in the context of ligand-bound signalling, we generated an AlphaFold model of the 2:2 ephrin-B2–EPHB2 extracellular complex and replaced the predicted EPHB2 ectodomains with the crystal structure (PDB 7S7K) aligned on the ligand-binding domains (Fig. 3a).

**Fig. 3:**
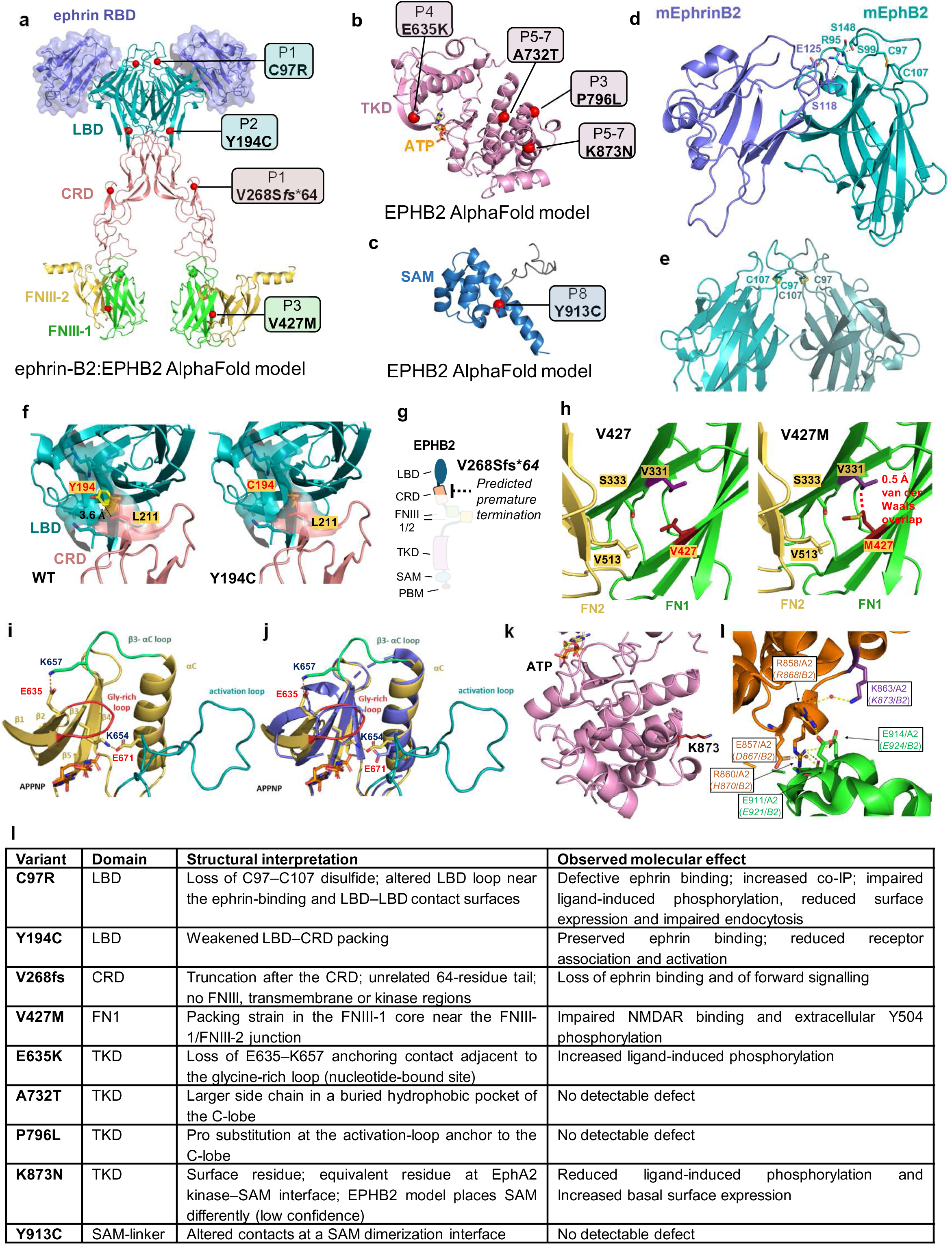
Structural analysis of EPHB2 variants implicates them in specific EPHB2 protein functions. **a**, Model of the 2:2 ephrin-B2 RBD–EPHB2 ectodomain complex (AlphaFold, with the EPHB2 ectodomains replaced by the crystal structure, PDB 7S7K, aligned on the LBDs). LBD, teal; CRD, salmon; FNIII-1, green; FNIII-2, gold; ephrin-B2 RBD, purple (transparent surface). Red spheres mark variant Cα positions, labelled by individual (P1–P8; Table 1). **b,c,** AlphaFold models of a single EPHB2 tyrosine kinase domain (TKD, pink; ATP, orange) (b) and SAM domain (blue) (c). P1, P3 and P5–P7 are siblings trans-heterozygous for two identical variants. **d,e,** C97 (P1) in the mouse EphB2–ephrin-B2 complex (PDB: 1KGY), showing the C97–C107 disulfide and nearby LBD residues (d) and the disulfides in the LBD–LBD arrangement of neighbouring receptors (e). **f,** Y194 (P2) packs against L211 at the LBD–CRD junction (WT, left; modelled Y194C, right). **g,** V268Sfs64 (P1) truncates the receptor after the CRD. **h,** V427 (P3) in FNIII-1 (WT, left; modelled V427M, right); the substitution creates a 0.5 Å van der Waals overlap with V331. **i,** E635 (P4) beside the glycine-rich loop of a nucleotide-bound (APPNP) Eph kinase. **j.** E635 in the EphA2 nucleotide-complex-based kinase model (PDB 5I9W; gold with structural features highlighted in Fig 3i) superimposed on apo EPHB2 (PDB 3ZFM; blue), where the glycine-rich loop and β3–αC loop are disordered. The E635–K657 contact and the K654–E671 salt bridge are labelled; APPNP, sticks. **k,** K873 (P5–P7) on the TKD surface. **l,** The equivalent residue at the EphA2 kinase–SAM interface. Hydrogen bonds are dashed lines. Residues are numbered as in human EPHB2 (NM_004442.7); equivalent residues in template structures are in parentheses. A732T, P796L and Y913C, which show no detectable functional defect, are in **Extended Data Fig. 3**. **m,** Table summarising the predicted impact of the variants on EPHB2 structure and observed molecular signalling consequences. Structural interpretations are based on experimental EPHB2 structures, AlphaFold models and, where indicated, homologous Eph receptor structures. Molecular effects summarize the biochemical and cell-biological assays presented in Fig. 2 and Extended Data Fig. 2.

Variants in the extracellular region are predicted to impair ligand recognition, receptor assembly or domain organisation. C97 forms an intramolecular disulfide bond with C107 that shapes the conformation of an LBD loop. This loop forms a hydrogen-bonding network that constrains R95, a key arginine contributing to ephrin recognition, and lies adjacent to residues involved in LBD-mediated receptor contacts (Fig. 3d,e). Disruption by C97R may therefore account for both its defective ephrin binding and altered receptor association, although the increased co-immunoprecipitation may represent aberrant rather than productive receptor assembly. Y194 is a conserved aromatic residue at the junction between LBD and cysteine-rich domain (CRD), where its side chain packs against L211 of the CRD and its hydroxyl group supports local hydrogen bonding in higher-resolution LBD structures (Fig. 3f). Y194C is predicted to weaken interdomain packing rather than directly alter the ephrin-binding surface, consistent with impaired receptor association and activation despite preserved ephrin recognition. The EPHB2 p.V268Sfs\**64* frameshift is predicted to cause premature termination in the CRD, replacing the remainder of the protein with an unrelated 64-residue tail and removing the downstream extracellular, transmembrane and intracellular regions, consistent with the observed loss of EphB2 protein function in the biochemical assays (Fig. 3g). The EPHB2 p.V427M variant is predicted to introduce local packing strain in FNIII-1 near the FNIII-1–FNIII-2 junction, potentially altering interdomain orientation and ectodomain conformation. This perturbation may contribute to reduced ephrin-B1-stimulated receptor activation and impaired EPHB2–NMDAR interaction, although the mechanistic links remain to be established (Fig. 3h).

Within the kinase domain, p.E635K alters a glutamate adjacent to the glycine-rich loop that contacts K657 in nucleotide- or inhibitor-bound Eph kinase structures, providing structural context for the increased ligand-induced tyrosine phosphorylation (Fig. 3i,j). p.K873N substitutes a solvent-exposed residue on the surface of the kinase domain (Fig. 3k). In EphA2 kinase–SAM structures, the equivalent residue makes an indirect, solvent-mediated contribution to the interdomain interface (Fig. 3l), raising the possibility that K873N alters kinase–SAM coupling, which is important for proper activation of tyrosine kinase^50^. However, the EPHB2 AlphaFold model places the SAM domain in a different orientation relative to the kinase domain than in EphA2 with low confidence (high inter-domain predicted aligned error; Extended Data Fig. 3a), providing uncertainty for structural interpretation. Three further variants are predicted to perturb local structure but showed no detectable defect in our assays (Extended Data Fig. 3b-e). p.A732T places a bulkier side chain in a buried hydrophobic pocket of the kinase C-lobe, p.P796L alters an invariant proline in the APE motif that anchors the end of the activation segment to the kinase core, and p.Y913C removes a residue buried in a SAM-domain dimerization interface observed in crystals. Together, the structure analyses combined with experimental observations indicate that the EPHB2 variants do not converge on a single molecular defect but disrupt specific steps of receptor signalling, although we did not find experimental evidence for some variants and request further verification.

### EphB2 expression correlates with callosal axon elongation

To understand how *EPHB2* variants might affect CC development, we studied EphB2 mRNA and protein expression in the developing mouse brain. Although EphB2 has been detected in mouse CPNs and surrounding tissues^26,27,30^, its spatiotemporal expression during CPN differentiation remains poorly defined. Mouse CPNs are born between embryonic day (E) 13 and E16/17 from radial glial cells (RGC), reaching a neurogenesis peak at E15.5 when >80% CPNs are born and completing their migration into superficial cortical layers by birth^51^. During this time, CPNs acquire a polarised morphology with basal side axons extending towards the callosal tract^52^. We thus isolated CPN neuron transcriptomic data from public embryonic mouse forebrain datasets and computed a *Ephb2* mRNA mean expression level over E13-E18 days^53^. Over this time, *Ephb2* mRNA levels increased, mirroring the expression of the callosal neuron transcription factor *Satb2* and other superficial CPN markers such as *Cux2* and *L1cam* (Fig. 4a and Extended Data Fig. 4a). A similar rise was observed for *Epha3* and *Epha4*, whose upregulation during CPN development was previously documented^54,55^, but not for *Ephb1* or *Ephb3*. In contrast, CPN expression of mRNAs encoding deep layer subcortical projection neuron markers such as *Fezf2* and *Bcl11b* or progenitor markers such as *Eomes* and *Pax6* decreased over the same period. Also, pooled forebrain neurons, neuroblasts, or radial glial cells showed a stable *Ephb2* mRNA expression level over the same period, suggesting that the increased expression of *Ephb2* mRNA in CNPs was cell-type specific.

**Fig. 4:**
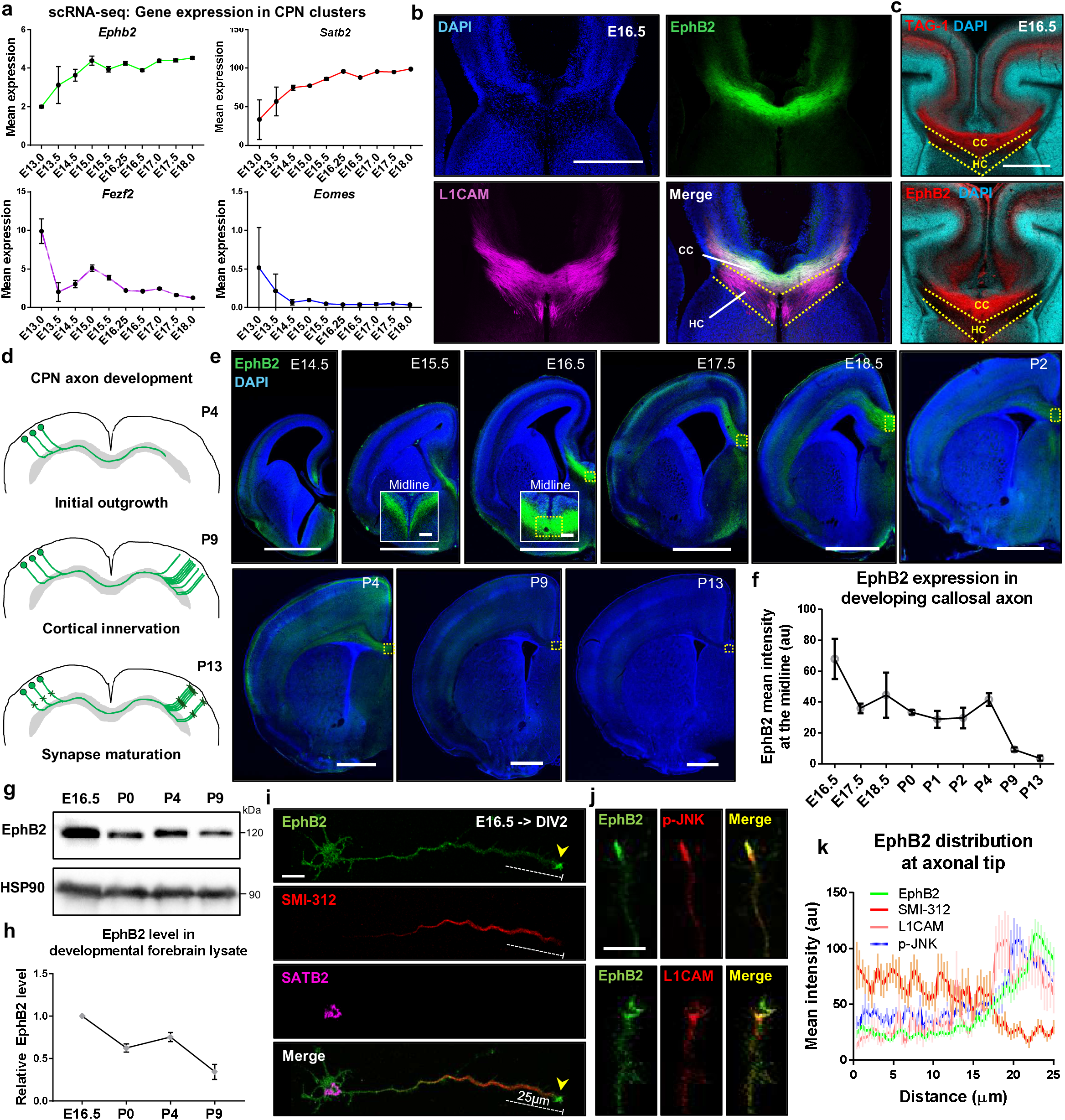
Spatially-restricted expression of EphB2 in CPN neurons at the time of callosal axon outgrowth. **a**, Mean CPN expression of *Ephb2*, *Satb2*, *Fezf2* and *Eomes* mRNA from E13-E18 mouse forebrain from single-cell RNA-sequencing datasets from La Manno et al., (2021), which includes 6,682 CPNs. **b,c,** EphB2 protein expression at the telencephalic midline in a coronal section of the E16.5 mouse forebrain. L1CAM (b) labels callosal axons (CC) and hippocampal commissural (HC) axons, whereas TAG-1 (c) labels only CC axons. Scale bar, 500 µm. **d,** Illustration of key stages of CPN axon development. **e,** EphB2 protein detection in coronal sections of ethe developing mouse forebrains at various embryonic ages. Scale bar, 1 mm, and 100 mm in insets. Solid line boxed insets show both sides of the midline, stippled boxes show unilateral signal at the midline quantified in f. **f,** Quantification of normalised EphB2 mean intensity at the midline, measured within the stippled rectangles in e. SEM; N=3. **g,** Detection of EphB2 and HSP90 in lysates from differently aged embryonic mouse forebrains. **h,** EphB2 expression levels in g normalised to HSP90. SEM; N=3. **i,** Expression of EphB2, SMI-312 and Satb2 in primary cortical neurons at DIV2. Arrowhead indicates the axonal terminus. Example of the distal-most 25 µm of the axon quantified in k are indicated by a stippled line. **j,** Images of growth cones from primary cortical neurons at DIV2 showing EphB2 localisation in comparison with phospho-JNK (p-JNK) and L1CAM. Scale bar, 10 µm. **k,** Mean expression levels of EphB2, SMI-312, L1CAM and p-JNK in the distal-most 25 mm of the axon. N=30 neurons for EphB2, 10 each for SMI-312, p-JNK, and L1CAM. Average values show as dark colours, and SEM of values are shown as light colours.

Next, we monitored the distribution of EphB2 protein in the developing mouse forebrain. At E16.5 when CPN pioneer axons enter the future CC, EphB2 was enriched at the leading edges of CPN axons (co-labelled with L1CAM/TAG-1) but not in hippocampal commissures (labelled with L1CAM only) located ventral to the CC (Fig. 4b,c). Subsequent profiling of EphB2 protein expression throughout CC development revealed its enrichment in CPN axons until postnatal day (P) 4 when CPN axons extend to the contralateral hemisphere but decrease in subsequent ages when axons enter cortical layers (P9) and establish synapses (P13), suggesting its potential involvement in CPN axon elongation (Fig. 4d-f). A similar trend was confirmed biochemically in developing forebrain lysates (Fig. 4g,h). Next, we examined the subcellular localisation of EphB2 in cultured cortical neurons. In CPNs identified via the expression of the Satb2 transcription factor, EphB2 levels appeared abundant at the axonal tip marked by phospho-JNK and L1CAM expression and absence of the axon shaft marker SMI-312 (Fig. 4i-k). We also examined protein distribution of other EPH receptors. Although other Eph receptors such as EphB1, EphA3, and EPHA4 were also expressed in the CC and cultured CPNs, EphA4 also showed enrichment at the tip of advancing axons (Extended Data Fig. 4b-e), suggesting that they may have a shared function in axon outgrowth^56^. Collectively, the temporally and spatially restricted expression of EPHB2 suggests that it may participate in callosal axon extension.

### Cell-autonomous function of EPHB2 in CPN development

Consistent with the link between our human *EPHB2* variants and CC dysgenesis, genetic studies of mouse Eph receptors revealed a prominent role of EPHB2 in CC development but do not define its specific functions in CPNs^26,28,29^. To address this, we used *in utero* electroporation (IUE) to introduce plasmids encoding wild type or EPHB2 variants, which showed deleterious molecular impacts (Fig. 2), into E15.5 mouse CPNs and examined CPN axon morphology at P4 (Fig. 5a,b). In all experiments we co-electroporated a plasmid encoding GFP that revealed CPN morphology and assumed that the endogenously expressed wild type mouse EphB2 would partially model the heterozygous state of the human *EPHB2* variants. Compared to vector control, expression of WT human EPHB2 did not result in a significantly different midline crossing rate or normalised axon volume but consistently showed a mild increase in CC width and CPN axon travel distance (Fig. 5c-f). In contrast, the KD mutant significantly decreased the CPN axon travel distance and mildly reduced CC width. The expression of C97R, V427M, and E635K variants did not result in any significant phenotypes, potentially due to their low expression in neurons (Fig. 2). Interestingly, the expression of V268Sfs from a proband with complete ACC, significantly impaired multiple aspects of CPN axon development. In contrast, expression of Y194C and K873N variants, from probands with callosal hyperplasia, showed increased axon travel distance compared to controls. We also observed that expression of WT, KD, Y194C, E635K, and K873N variants disrupted apical migration of CPNs that could potentially affect local connectivity within the cortex of affected individuals (Extended Data Fig. 5a,b), although laminar patterning and functional cortical connectivity in the affected individuals are unknown. Together, these data provide evidence that EPHB2 and its variants can affect CPN axon development and neuronal migration in a cell-autonomous manner.

**Fig. 5:**
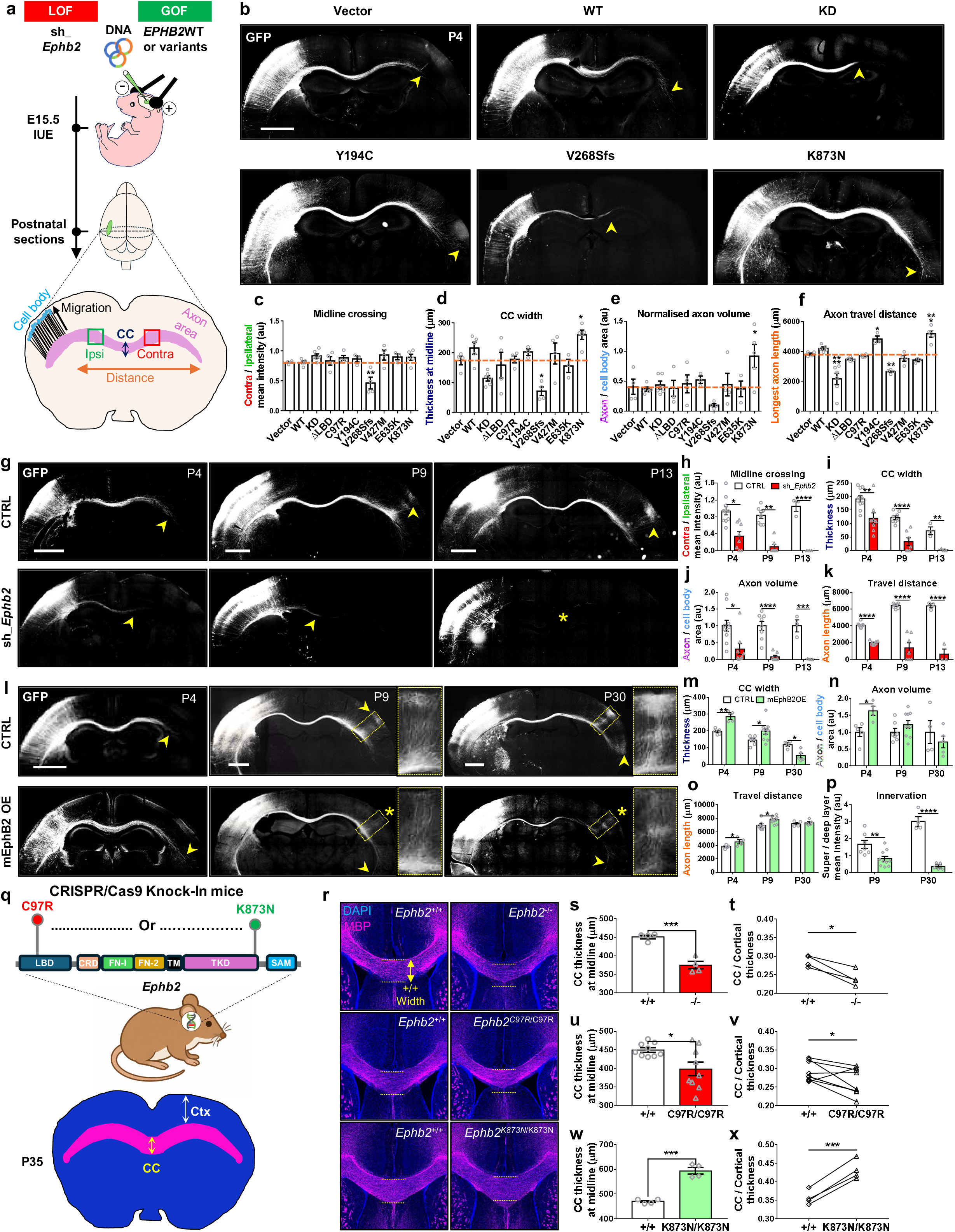
EphB2 functions cell-autonomously to regulate callosal development. **a**, Schematic of the *in utero* electroporation (IUE) workflow. Plasmids were unilaterally electroporated into the cortical ventricular zone at E15.5. Postnatal mouse brains were dissected and analysed. Midline crossing rate of CPN axons was obtained by dividing the mean intensity of contralateral CPN axons (red box) by the mean intensity ipsilateral CPN axons (green box). CC width was calculated as the distance between dorsal-most and ventral-most axons at the CC midline. Normalised axon volume was measured by dividing axon area (pink) by cell body area (light blue). Axon travel distance was measured as the length of the fifth-longest axon. **b,** P4 coronal brain sections following IUE of expression plasmids encoding variants (or empty expression plasmids) and a GFP expression plasmid. Arrowheads indicate the terminus of longest axons. Scale bar, 1 mm. **c-f,** Quantification of midline crossing rate (c), CC width (d), normalised axon volume (e), and axon travel distance (f) in P4 coronal brain sections following IUE. Orange dashed lines indicate the average values in the empty expression plasmid controls. SEM; N = 3-6. **g,** Images of P4, P9, and P13 coronal brain sections following IUE of control (CTRL) or sh_*Ephb2* plasmids expressing EphB2-targetting shRNA. Scale bar, 1 mm. Arrowheads indicate the terminus of the longest axons. **h-k,** Quantification of midline crossing rate (h), CC width (i), normalised axon volume (j), and axon travel distance (k) in P4, P9, and P13 brain sections following control or sh_*Ephb2* plasmid IUE. SEM; N = 8-10 (P4), 7-8 (P9), 3 (P13). **l,** Images of P4, P9, and P30 coronal brain sections after control (CTRL) or mEphB2 expression plasmid IUE. Scale bar, 1 mm. Arrowheads indicate the terminus of the longest axons. Stippled boxes highlight axon termination and cortical neuron innervation zones in the contralateral cortices. **m-p,** Quantification of CC width (m), normalised axon volume (n), axon travel distance (o), and innervation rate (p) in P4, P9, and P30 coronal brain sections following CTRL or mEphB2 IUE. SEM; N = 4-5 (P4), 7-10 (P9), 4-5 (P30). **q,** Location of quantified cortical (Ctx) and callosal (CC) thickness in mice homozygous for EphB2 knock-out, human variant alleles and littermate control mice. **r,** Midline images from P35 mouse coronal brain sections of homozygous *EphB2* mutant or control mice. **s,u,w,** Absolute CC thickness expressed as mm. N=4 for *Ephb2^-/-^*, 9 for *Ephb2^C97R/C97R^*, 4 for *Ephb2^K^*^873^*^N/K^*^873^*^N^* mice, with an equal number of littermate controls for each genotype. **t,v,x,** CC thickness normalised to cortical thickness. Littermates were used for all analyses.

### EphB2 regulates callosal axon growth in a stage- and dosage-dependent manner

Next, to examine the effects of loss of endogenous EphB2, we used IUE of shRNAs that specifically and efficiently reduce EphB2 protein levels (sh_*Ephb2*) in combination with a GFP expression plasmid at E15.5 (Extended Data Fig. 5c-f)^57,58^. First, we monitored early CPN developmental processes, including cortical progenitor proliferation, survival, apical migration, and CPN cell fate specification. The fraction of electroporated neurons that co-expressed the KI-67 proliferation marker, or the cleaved caspase-3 apoptosis marker was unchanged compared to controls at P0 (Extended Data Fig. 5g-j) and subsequent ages (data not shown). Interestingly, we were unable to observe migration defects in EphB2 depletion (Extended Data Fig. 5k,l). Consistent with the intact cell positioning, neurons electroporated with controls or sh_*Ephb2* expressed the superficial cortical layer (L2-4)-specific transcription factor Lhx2 and did not express deep cortical layer specific transcription factors, CTIP2 (L5) or TBR1 (L6), while the fraction of neurons expressing the CPN-specific transcription factor SATB2 was unchanged (Extended Data Fig. 5m-o). These data show that EphB2 depletion does not substantially affect proliferation, cell-type specification, migration, or survival, suggesting a less prominent role of endogenous EphB2 in these processes.

Next, we assessed the effect of EphB2 loss on different steps of CPN axon development (Fig. 4d). Compared to controls, EphB2 depletion resulted in decreased midline crossing, CC width, axon volume and axon travel distance at P4 (Fig. 5g-k). Whole mount tissue clearing of P4 brains did not reveal any evidence that EphB2 depletion caused a redirection of CPN axons from the CC into another major axon tracts (Supplementary Videos 1 and 2). At P9 and P13, EphB2 depletion resulted in an almost complete absence of CPN axons from the CC. Importantly, co-electroporation of the shRNA with a plasmid encoding an *EPHB2* mRNA that is resistant to the shRNA (EPHB2res) partially restored midline crossing, axon volume, and travel distance compared to shRNA alone (Extended Data Fig. 5p-s). Together, these data suggest that EphB2 play an important role in CPN axon extension.

In contrast, overexpression of mouse EphB2 (mEphB2) resulted in robust enhancement of initial CPN axon outgrowth; increased CC width, axon volume, and axon travel distance without affecting midline crossing at P4 (Fig. 5l-o and Extended Data Fig. 5t). However, maintained mEphB2 expression prompted axon traversal within the callosal tract at P9 and prevented proper innervation and arborisation of axons (Fig. 5p), resulting in a decrease in the final number of axons at P30. Interestingly, expression of K873N variant also showed decrease in axon volume similar to wild type, whereas CPN axons expressing the Y194C variant maintained axon volume but retained exuberant projections in deeper cortical layers (Extended Data Fig. 5u). These observations highlight the importance of tightly regulated EphB2 expression during CPN axon outgrowth stage (Fig. 4e,f), as well as the requirement for intact EphB2 molecular functions for proper CC development and morphology. Together, these findings support a stage- and dosage-dependent role for EphB2 in CPN development: CPN-intrinsic EphB2 promotes interhemispheric axon extension, whereas its dysregulated or sustained expression perturbs correct local circuit formation within cortices.

### Mouse models of human *EPHB2* variants recapitulate their callosal phenotypes

To determine whether the EPHB2 variants identified in affected probands are sufficient to alter CC development, we generated mice homozygous for the p.C97R or p.K873N variants (Fig. 5q), associated with the most striking morphological defects in the anterior CC of probands 1 and 5-7, respectively (Fig. 1e). We then compared their CC thickness to littermate controls and mice homozygous for the *EphB2^-^*knockout allele^26,29^. Compared to controls, *EphB2^C97R^*^/*C97R*^ mice displayed a similar degree of CC reduction as *EphB2^-/-^* mice, suggesting the p.C97R variant is a loss of function (Fig. 5r-v). On the other hand, *EphB2^K^*^873^*^N/K^*^873^*^N^* mice showed a significant increase in CC thickness suggesting that the p.K873N variant behaves as a gain of function (Fig. 5w,x). Together, these data establish mouse models of p.C97R or p.K873N EPHB2 human variants and align with their biochemical and cell biological analyses.

### EphB2 promotes callosal axon elongation through spatial regulation of active myosin and F-actin at growth cones

Ephrin-mediated Eph receptor signalling is known to promote growth cone collapse and axon retraction^25^. However, EPHB2 variant carrier CC phenotypes and the elevated and spatially restricted EphB2 expression at the time of CPN growth strongly suggest a growth promoting role of EphB2. Thus, we next asked whether and how EphB2 can promote axon elongation in individual neurons. To study the cellular mechanism of EphB2 function, we co-cultured age-matched control and EphB2-depleted mouse CPNs, expressing GFP or RFP, respectively (Fig. 6a). Neurite lengths were analysed at three, five or eight days in vitro (DIV), corresponding to immature neurite formation, initial axon elongation, and the completion of axon elongation stages, respectively^52,59^. At DIV3, no difference was observed in average neurite length between control and shRNA neurons (Fig. 6b,c). However, at DIV5 when one of these neurites became an extending axon, its length was significantly reduced in neurons with EphB2 depletion compared to control neurons. By DIV8, this difference became even more dramatic. Consistent with this, we observed that EphB2 expression was elevated at DIV5, which was effectively inhibited by sh_*Ephb2* (Extended Data Fig. 6a-d). Cell fate acquisition and axon specification was unaffected (Extended Data Fig. 6e-h), suggesting a selective role of EphB2 in axon elongation. Interestingly, treatment with clustered-ephrin-B1 or Fc, did not change axon length in control or sh_*Ephb2-*treated neurons (Fig. 6c and Extended Data Fig. 6i), suggesting a ligand-independent role of EphB2 in axon elongation.

**Fig. 6:**
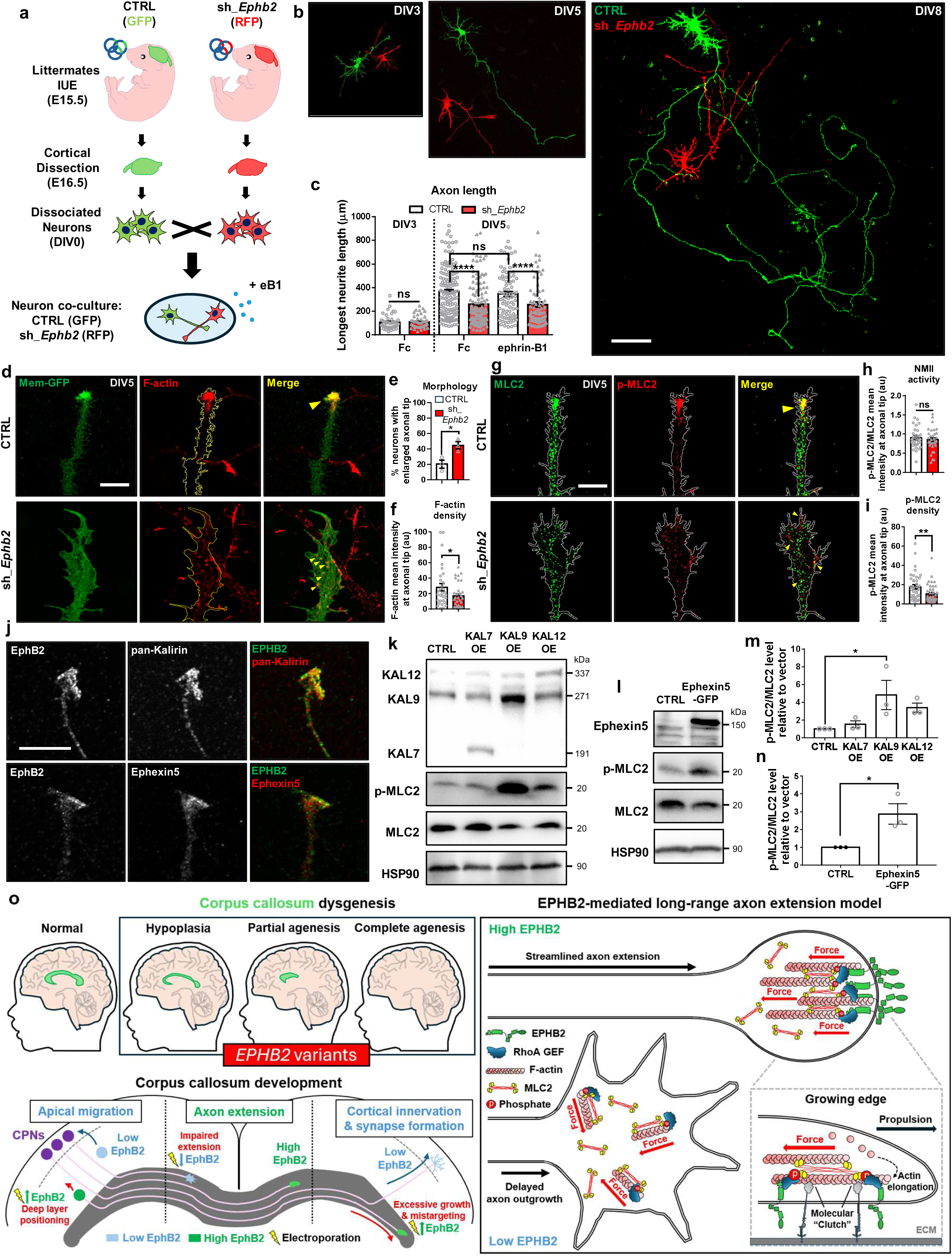
EphB2 promotes callosal axon outgrowth through spatial restriction of actin-myosin at the growth cone. **a**, The workflow of CPN neuron co-culture assay. IUE electroporation of control or sh_*Ephb2* plasmids and, respectively, GFP and RFP expression plasmids was carried out at in separate cohorts at E15.5. A day later, cortices were dissection and cortical neurons dissociated. A mixture of control or sh_*Ephb2* plasmid transfected neurons was co-cultured and treated with Fc IgG fragment, pre-clustered ephrin-B1. **b,** Co-cultured control plasmid (green) or sh_*Ephb2* plasmid (red) cortical neurons at various days of culture in vitro (DIV). Scale bar, 100 µm. **c,** Quantification of the longest neurite length in electroporated neurons at DIV5 grown in presence of either Fc IgG fragment or pre-clustered ephrin-B1. SEM; N>100 neurons from 4 independent experiments. **d,** GFP expression and F-actin localisation revealed with phalloidin in axonal growth cones of DIV5 primary cortical neurons electroporated with either control or sh_*Ephb2* plasmids. Arrowheads indicate F-actin particles. Scale bar, 10 µm. **e,** Proportion of primary neurons with enlarged axonal terminus (diameter > 100 μm^2^) following electroporation with either CTRL or sh_*Ephb2*. SEM; N=3 (>30 neurons in each replicate). **f,** Mean actin intensity in axonal termini of DIV5 primary neurons electroporated with either control or sh_*Ephb2* plasmids. SEM; N>30 neurons from 3 independent experiments. **g,** Localisation of MLC2 and p-MLC2 in axonal growth cones of DIV5 primary cortical neurons electroporated with either control or sh_*Ephb2* plasmids. Arrowheads indicate p-MLC2/MLC2 co-localisation. Scale bar, 10 µm. **h,** NMII activity expressed as the ratio of mean p-MLC2 to MLC2 intensities at axonal termini of primary neurons electroporated with either control or sh_*Ephb2* plasmids. SEM; N>30 neurons from 3 independent experiments. **i,** Mean p-MLC2 intensity in axonal termini of primary neurons electroporated with either control or sh_*Ephb2* plasmids. SEM; N>40 neurons from 3 independent experiments. **j,** Kalirin (all isoforms, top) or Ephexin5 (bottom), EphB2 expression in axonal growth cones from DIV2 primary cortical neurons expressing the CPN marker SATB2. Scale bar, 10 µm. **k,l,** Immunoblots of cell lysates from DIV6 rat primary cortical neurons following nucleofection with plasmids encoding individual Kalirin isoforms (k) or Ephexin5-GFP (l). **m,n,** Quantification of p-MLC2/MLC2 levels in cell lysates from DIV6 rat primary cortical neurons following overexpression of individual Kalirin isoforms (m) or Ephexin5-GFP (n), normalised to vector controls. SEM; N=3. **o,** Diagrams illustrating our interpretation of the impact of human *EPHB2* variants on corpus callosum morphology (top left), the role of EphB2 in different stages of CPN development (bottom left), and the molecular and cellular mechanisms underlying EphB2’s function in axonal growth (right).

Growth cones of actively extending axons typically display a streamlined, highly polarized morphology driven by a robust F-actin-assembly, whereas pausing or slow-moving growth cones transition into an expanded, fan-shaped architecture optimized for environmental exploration^60–63^. At DIV5, when a majority of control CPNs exhibited an extending morphology, EphB2 depletion promoted large, fan-shaped morphology at the axonal terminus (Fig. 6d,e and Extended Data Fig.6j). Furthermore, while control axons showed high F-actin levels concentrated at their tips, neurons with diminished EphB2 levels showed lower levels and diffuse distribution of F-actin within the axonal growth cones (Fig. 6f).

Phosphorylated active non-muscle myosin II (NMII) in neuronal growth cones organises the spatial distribution of F-actin and regulates axon outgrowth in cortical neurons^35^. We therefore examined the subcellular co-localisation of EphB2 and phosphorylated-myosin light chain 2 (p-MLC2) in native CPNs and found that, in streamlined growth cones, p-MLC2 localised within EphB2-enriched domains (Extended Data Fig. 6k). In contrast, fan-shaped growth cones showed diffuse EphB2 and p-MLC2 localisation with little spatial overlap. Interestingly, EphB2 depletion led to a diffuse distribution of p-MLC2 in CPN axonal tips, without affecting total levels or activity (p-MLC2/MLC2) (Fig. 6g-i and Extended Data Fig. 6l,m), suggesting that EphB2 promotes a local enrichment of active NMII within actively extending CPN axon growth cones.

To identify the specific cytoskeletal processes disrupted by EphB2 depletion, we compared the axonal length of DIV5 control and EphB2-depleted cortical neurons, in which microtubule or actin polymerisation, myosin contractility, or actin nucleation and branching were inhibited at DIV4 (Extended Data Fig. 6n,o). Compared to DMSO vehicle treatment, blocking microtubule assembly with nocodazole severely reduced axon outgrowth both in control and EphB2-depleted neurons, confirming the primary involvement of microtubules in axon extension. Interestingly, disruption in actin polymerization with Latrunculin A treatment also significantly inhibited axon outgrowth as well as promoting fan-shaped growth cone morphology, similar to shRNA alone. Furthermore, Latrunculin A treatment of EphB2-depleted neurons did not further decrease axon length, suggesting that both manipulations may target the same cytoskeletal pathway. Next, treatment with either CK-666, an inhibitor of arp2/3-mediated actin nucleation and branching, or blebbistatin, an inhibitor blocking myosin-mediated F-actin contraction, promoted the formation of extending growth cone morphologies and axon outgrowth in control neurons compared to DMSO treatment. Strikingly, both compounds restored axon lengths as well as the streamlined growth cone morphology in EphB2-depleted neurons relative to controls. Together, these data argue that EphB2 promotes axon outgrowth in CPNs by modulating myosin-actin dynamics.

To identify candidate regulators linking EphB2 to actomyosin organisation, we examined the distribution of the Rho-family guanine nucleotide exchange factors (Rho GEFs) Kalirin and Ephexin5, both previously implicated as effectors of EphB-regulated cytoskeletal dynamics in developing neurons^64,65^. In DIV5 cortical neurons, antibodies detecting multiple isoforms of Kalirin and Ephexin5 revealed their presence within EphB2-rich domains at axonal termini (Fig. 6j), suggesting that these proteins may participate in EphB2-dependent regulation of the growth cone cytoskeleton. Because EphB2 depletion decreased p-MLC2 localisation to axonal tips (Fig. 6g-i), we next asked whether Kalirins or Ephexin5 can regulate NMII activation. Overexpression of the long Kalirin isoforms Kalirin9 and Kalirin12 in cortical neurons increased the p-MLC2/MLC2 ratio compared to control neurons, whereas Kalirin7 had little effect (Fig. 6k,m). Similarly, Ephexin5 overexpression increased p-MLC2 levels without altering total MLC2 abundance (Fig. 6l,n). Together, these findings identify RhoA-specific GEFs, long Kalirins and Ephexin5, as candidate mediators linking EphB2 signalling to activation of the actomyosin machinery in developing CPNs to mediate long-range axon outgrowth.

## Discussion

Cortical expansion during brain evolution coincided with a structural increase in interhemispheric connectivity through the CC^1^. Long-range axon extension is fundamental to CC development and the assembly of many other neural circuits^9^, yet the underlying mechanisms remain incompletely understood. Here, we show that rare variants in *EPHB2* coding sequence are associated with distinct CC dysgenesis. Structural and biochemical analyses reveal variant-specific molecular effects on different aspects of EPH signalling, whereas manipulation of EphB2 in developing mouse CPNs uncovers a stage- and dosage-dependent role in CPN axon development. Moreover, we demonstrate that EphB2 promotes CPN axon elongation through spatial organisation of actomyosin at axonal growth cones. Together, our findings link human *EPHB2* variation to a spectrum of callosal abnormalities and reveal a previously unrecognised role for EphB2 in promoting long-range axon extension (Fig. 6o).

Few recent studies have investigated and identified functional defects in one *EPHB2* variant in mice, revealing abnormalities in neural circuit activity and behavioural phenotypes, including autism-like behaviours and anxiety^22,23^. However, anatomical abnormalities in affected individuals that may underlie these circuit-level and behavioural deficits remain poorly understood. Our study demonstrates that rare *EPHB2* variants are associated with a spectrum of CC abnormalities in humans. CPN-specific expression of human EPHB2 variants and knock-in mouse models revealed partial correspondence between the direction of the CC phenotypes in mice and the callosal abnormalities observed in the respective individuals, providing evidence for a causal link between *EPHB2* variation and CC dysgenesis. Beyond CC dysgenesis, affected individuals also exhibited defects in other known roles of EphB2 in neural circuit assembly including visual defects and anxiety^66,67^. Notably, a variant identified in an individual with chronic anxiety selectively disrupted EPHB2-NMDAR interaction, a complex implicated in amygdala-dependent anxiety, pain, NMDAR-encephalitis, and Alzheimer’s Disease^67–70^. This finding provides additional links between *EPHB2* variants and the clinical manifestations observed in affected individuals, warranting further investigation.

Various Eph receptors are expressed in CPNs and their surrounding tissues^26,27^. However, the temporal dynamics of Eph receptor expression and the functional contributions of individual receptors remain incompletely understood, complicating isolation of their stage-specific roles in CPN development. Early studies of EphB2 in CC development highlighted its role in midline structures for guiding CPN axons across the interhemispheric midline^26,30^, whereas subsequent studies provided evidence for additional CPN-intrinsic functions^28,29^. Here, we show that EphB2 expression increases in CPN axons concomitantly with the onset of axon elongation and subsequently declines during later processes. CPN-specific gain- and loss-of-function experiments demonstrate that EphB2 promotes interhemispheric axon extension, whereas its dysregulated or sustained expression perturbs local circuit connectivity within cortices. These findings expand our understanding of EphB2 function in CC development, from its established role in midline structures for axon guidance to a cell-autonomous function in promoting CPN axon outgrowth.

One mechanism that promotes rapid axon elongation involves the formation of molecular clutches that anchor F-actin to extracellular matrix proteins, thereby converting NMII-driven retrograde actin flow into forward traction^71^. We propose that EphB2 organises active NMII into a compact architecture at axonal tips, potentially aligning the direction of NMII-generated forces to facilitate coordinated F-actin dynamics and promote efficient axon extension. Consistent with this model, EphB2 has been associated with L1CAM and focal adhesion kinase^72,73^, both of which are crucial for molecular clutch function. We further identified the candidate effectors Kalirin9/12 and Ephexin5, EPHB2-interacting RhoA GEFs^64,65^, linking active NMII at axonal growth cones. This is consistent with previous observations for Kalirin long isoforms in promoting axon elongation^74,75^. These results provide evidence of a previously unrecognised mode of Eph signalling whereby EphB2 promotes long-range axon elongation, expanding the functional repertoire of Eph receptors. More broadly, our study links human genetic variants to individualised abnormalities in brain connectivity, providing a framework for understanding how heterogeneous circuit-level phenotypes arise in neurodevelopmental disorders.

## Methods

### -Identification of *EPHB2* variants

The individuals were identified through GeneMatcher^76^ or the International Research Consortium for the Corpus Callosum and Cerebral Connectivity (IRC5). All *EPHB2* variants were identified by whole exome sequencing (WES) performed on whole blood DNA- from the individuals. The variants were defined based on standardised nomenclature from the reference human genome GRCh38 (hg38) and EPHB2 isoform 2 transcript (NM_004442.7) following its expression abundancy in human tissues according to the Genotype-Tissue Expression (GTEx) data platform. The allele frequency of each variant was determined from genomic sequencing data derived from gnomAD^77^.

### -*EPHB2* rare variant association analysis using cohort in UKB with Corpus Callosum MRI scans

Brain MRI data were obtained from the UK Biobank (UKB) imaging study. T1-weighted structural images were acquired at 1 mm isotropic resolution using a 3D MPRAGE sequence on 3T Siemens Skyra scanners with a 32-channel receive head coil^78^. Images were processed through the UKB automated brain imaging pipeline, which includes gradient distortion correction, brain extraction, and registration to standard space^79^. Subcortical volumetric segmentation was performed using FreeSurfer’s automated segmentation tool (aseg), which assigns neuroanatomical labels to each voxel based on a probabilistic atlas^80^. The corpus callosum (CC) was further parcellated into five subregions using FreeSurfer’s mri_cc tool, which segments the CC within the aseg volume into equally spaced divisions along its anterior-posterior axis with a default lateral thickness of 5 mm. The five resulting subregions are: CC-Anterior (genu; UKB Data-Field 26535), CC-Mid-Anterior (Data-Field 26534), CC-Central (Data-Field 26533), CC-Mid-Posterior (Data-Field 26532), and CC-Posterior (splenium; Data-Field 26531). All volumes are reported in mm³. We only focused on the data in the first imaging visit (instance 2). An average CC volume was calculated among those five subregions and used as the primary phenotype. This composite measure was chosen for several reasons. First, averaging across subregions provides a more robust estimate of overall CC morphology by reducing measurement noise associated with any single subregion. Second, a single summary measure maximises statistical power in genetic analyses by limiting the multiple-testing burden that would arise from analysing each subregion independently. Third, genetic variants influencing general white matter tract development or myelination are expected to exert broadly distributed effects across the CC, making a global measure well suited to capture such shared variance. Subregion-specific analyses may be pursued in follow-up to characterise any regionally selective genetic effects.

Confound variables for brain imaging analyses were selected in accordance with recommendations from the UKB imaging team^81^. These included age at imaging, age², sex, age * sex, head motion (UKB Data-Fields 25741–25742), scanner table position (X, Y, and Z coordinates; UKB Data-Fields 25756–25758), imaging assessment centre, and scan-date-related temporal drifts. 10 genetic principal components were additionally included to control for population stratification.

Whole genome sequencing (WGS) data for approximately 500,000 UKB participants were generated using Illumina NovaSeq 6000 sequencing machines at an average coverage of 32.5×^82^. Sequence reads were mapped to the GRCh38 human reference genome using BWA. Single-nucleotide polymorphisms (SNPs) and short insertions/deletions (indels) were jointly called across all participants using GraphTyper and GATK haplotypeCaller^83,84^. The jointly called variant data produced were used for downstream analyses (UKB Data-Field 23374). Variants within the EPHB2 gene were extracted from the WGS data based on the genomic coordinates chr1:22,710,839–22,921,500 (GRCh38). Individual-level quality control filters were applied, excluding genotypes with read depth (DP) < 25 or genotype quality (GQ) < 25. For the final burden analysis, only rare variants with a minor allele frequency (MAF) < 0.01 and a minimum minor allele count (MAC) > 5 were retained. Functional annotation of variants was performed using ANNOVAR^85^. Nonsynonymous (nonsyn) and noncoding groups of variants were identified for burden analyses. Participants were excluded based on several criteria designed to minimise confounding of corpus callosum volumetric and genetic analyses. To address population stratification in genetic analyses, the sample was restricted to individuals of European ancestry, and participants whose genetic sex did not match self-reported sex were excluded to ensure data integrity. Relatedness closer than cousin was used as an additional exclusion criterion to prevent inflation of test statistics due to cryptic familial structure. Individuals with any neurological conditions including degenerative, vascular, traumatic, and infectious brain pathologies were excluded, may independently affect corpus callosum volume, potentially confounding genetic association analyses (Supplementary Table 2). Analyses were conducted in two exclusive cohorts: individuals without any mental health conditions and individuals with mental health conditions. The mental health conditions include autism spectrum disorder (ASD), attention deficit and hyperactivity disorders (ADHD), depression, schizophrenia, anxiety, and others (UKB Data-Field 29000). The inclusion of mental health conditions was motivated by prior evidence that EPHB2 genetic variants were associated with autistic individuals^21^ and caused behavioural deficits in animals^22,23^. Furthermore, all eight patients in our cohort showing corpus callosum dysgenesis also exhibit diverse comorbid mental health conditions (Extended Data Fig. 1b)

Gene-based rare variant association analysis was performed using the Sequence Kernel Association Test (SKAT) and Burden test as implemented in the SKAT R package^86^. SKAT was used as the primary gene-based rare variant association test. The Burden test was additionally applied as a sensitivity analysis to evaluate whether rare variants within each gene act in a uniform direction. A Bonferroni correction was applied to the primary SKAT results only. Rare variants were defined as those with a MAF < 0.01. EPHB2 variants were grouped into noncoding and nonsyn sets, and the SKAT test was applied to assess the joint association of rare variants within EPHB2 with mean CC volume. The imaging confounding variables described above, together with 10 genetic principal components, were included as covariates in all models.

### -Mouse lines and animal care

Experiments described were conducted following the guidelines for animal care of the Animal Care Commitee of the Institut de Recherches Cliniques de Montréal (protocol 2023-11 AK). All mutant mouse (*Mus musculus*) lines were maintained in the C57BL/6J background by regular backcrossing to the C57BL/6J line (stock number 000664; The Jackson Laboratory) otherwise indicated. Pregnant CD1 and B6D2F1 females were used for in utero electroporation experiments to avoid problems with cannibalism in C57BL/6J strain after surgery. All tested phenotypes for in utero electroporation experiments were reproducible in both CD1 and B6D2F1 strains. All mice were housed at around 22-23 °C on a 12-h-light/12-h-dark cycle and fed ad libitum regular chow and water.

### -Plasmids

The human EPHB2 cDNA was synthesised and subcloned into both pCDNA3.1 (+) and pCAGGs-ES vector with or without C-terminal tag (GFP or Myc) (Genscript). All EPHB2 constructs harbouring point variants including all human variants and annotated controls (kinase-dead, ΔLBD, and rescue constructs) used in the study were generated using site-directed mutagenesis (Genscript).

The following plasmids were purchased from indicated suppliers: pCAG-mouseEphb2 and pCAG-myrRFP (Addgene); pLKO-sh_Ephb2 and pLKO-sh_Random (Millipore Sigma). The following plasmids are generous gifts from indicated research groups: pSUPER-sh_Ephb2 and pSUPER-vector (Dr. Dalva); pSUPER-shScramble, pSUPER-sh_Ephexin5, pEF-Ephexin5wt/LQR/Y361F, and pGFP-E5-GFP (Dr. Margolis); pEAK10-His-Myc-Kalirin7, pEAK10-His-Myc-Kalirin9, and pEAK10-His-Myc-Kalirin12 (Drs. Penzes and Parnell). All plasmids were repeatedly verified by full-length sequencing or by Sanger sequencing.

### -Generation of EPHB2 expression constructs with human variants

The human *EPHB2* cDNA was synthesized and subcloned into both pCDNA3.1 (+) and pCAGGs-ES vector with or without C-terminal tag (GFP or Myc) (Genscript). All EPHB2 constructs harbouring point variants including all human variants and annotated controls (kinase-dead, ΔLBD, and rescue constructs) used in the study were generated using site-directed mutagenesis (Genscript).

### -Plasmids

pCAG-mouseEPHB2, pCAG-myrRFP, pSuper-shScramble plasmids were purchased from Addgene. pSuper-sh_*Ephb2* and pSuper-vector plasmids are gifts from Dr. Dalva. pLKO-sh_*Ephb2* and pLKO-sh_*Random* were purchased from Sigma. All plasmids were verified by full-length sequencing (Genscript) after generation and repeatedly checked by Sanger sequencing using internal primers after plasmid amplification using maxi-prep kit (Qiagen).

### -Antibodies and chemicals

The following antibodies were purchased from indicated suppliers: goat anti-EPHB2 (WB/ICC/IHC 1:500, AF467), mouse anti-EPHB3 (WB 1:1,000, IHC 1:500, ICC 1:100, MAP5667), goat anti-Contactin2/TAG-1 (IHC 1:1,000, AF4439) (R&D Systems); rabbit anti-EPHB1 (Q-20) (WB 1:1,000, ICC/IHC 1:100, sc-926), rabbit anti-EPHA3 (L-18) (WB 1:1,000, ICC/IHC 1:100, sc-920), mouse anti-phospho-tyrosine (pY20) (WB 1:1000, sc-508), mouse anti-Myc (9E10) (WB 1:1000, ICC 1:500, sc-40) (Santa Cruz Biotechnology); mouse anti-EPHA4 (WB 1:1,000, ICC/IHC 1:100, 37-1600), rabbit anti-GFP (WB 1:10,000, ICC/IHC 1:1,000, A-11122), rabbit anti-MYL9 (ICC 1:100, PA517624), mouse anti-phospho-MYL9 (Ser19) (ICC 1:100, MA5-15163), DAPI (ICC/IHC 1:500, D1306) (Thermo Fisher Scientific); goat anti-GFP (ICC 1:1,000, ab6673), mouse anti-SATB2 (ICC/IHC 1:1,000, ab51502), rabbit anti-LHX2 (IHC 1:500, ab184337), rabbit anti-TBR1 (IHC 1:500, ab31940), rabbit anti-KI-67 (IHC 1:500, ab16667), phalloidin-iFluor 633 (1:1,000, ab176758) (abcam); rabbit anti-HSP90 (WB 1:1,000, 4874), rabbit anti-phospho-JNK (WB 1:1,000, ICC 1:100, 9255), rabbit anti-phospho-ERK (WB 1:1,000, ICC 1:100, 9101), rabbit anti-Cleaved Caspase-3 (IHC 1:500, 9661), rabbit anti-Myosin Light Chain 2 (WB 1:1,000, 8505), rabbit anti-phospho-Myosin Light Chain 2 (Ser19) (WB 1:1,000, 3671) (Cell Signaling Technology); mouse anti-ACTB (WB 1:10,000, A5441), rat anti-Neural Cell Adhesion Molecule L1 (L1CAM) (ICC/IHC 1:500, MAB5272), rat anti-CTIP2/BCL11B (IHC 1:500, MABE1045), Nocodazole (M1404), Latrunculin A (L5163), (-)-Blebbistatin (B0560) (Millipore Sigma); chicken anti-GFP (ICC 1:1,000, GFP-1010), chicken anti-Myelin Basic Protein (IHC 1:500, MBP) (Aves Labs); mouse anti-Neurofilament Marker (SMI-312) (ICC 1:500, 837904, Biolegend); rat anti-RFP (ICC/IHC 1:1,000, 5f8, Chromotek); rabbit anti-RFP (WB/IHC 1:1,000, 600-401-379, Rockland); CK-666 (HY-16926, MedChemExpress)

The following antibodies are generous gifts from indicated research groups: rabbit anti-N-Ephexin5 (WB 1:1,000) (Dr. Greenberg); rabbit anti-Ephexin5 (ICC 1:100) (Dr. Zito); rabbit anti-pan-Kalirin (WB 1:1000, ICC 1:100) (Dr. Penzes).

### -Immunocytochemistry

Neuronal cultures and HeLa cells were fixed by directly adding 4% paraformaldehyde (PFA) (with 6% sucrose) in PBS (1x) into the cell-containing media for 12 minutes at room temperature (RT). After fixation, the cells were rinsed with PBS three times and permeabilized with 0.2% PBST (0.2% Triton X-100 (Millipore Sigma 108603) in 1x PBS) for 5 minutes. Cells were rinsed with PBS three times and incubated with blocking buffer (10% inactivated horse serum (IHS) in PBS) for 1hr. Then, the cells were incubated with primary antibodies in blocking buffer overnight at 4 °C. Next day, the cells were rinsed with PBS three times and incubated with secondary antibodies in blocking buffer for 3hrs at RT. For every experiment, DAPI was added during secondary antibody incubation at a 1:500 dilution for nuclei staining. Following three times rinse with PBS, the cells were mounted using Mowiol solution (Millipore Sigma 81381).

### -Immunohistochemistry

Brain tissues from P14 and older mice were fixed by sequential transcardial perfusion with PBS and 4% PFA followed by overnight fixation in 4% PFA at 4 °C with shaking. Brain tissues of postnatal mice younger than P14 were fixed by transcardial perfusion directly with 4% PFA and post-fixed with 4% PFA overnight. Embryonic mouse brains were first dissected out from skull and fixed in 4% PFA 3-5hrs. After fixation, brains were washed with PBS three times for 10 minutes each and kept in PBS added with 0.02% sodium azide until use. Before cutting brains, the brain tissues were embedded in melted 4% agarose. Once the agarose was consolidated, the agarose blocks including brain tissues were directed for sectioning using vibratome (Leica VT1200). All tissue sections were cut at 70μm and collected in PBS added with 0.02% sodium azide. For immunofluorescence staining, sections were first incubated in blocking buffer (5% IHS and 2% bovine serum albumin (BSA) in 0.2% PBST) for 1hr at RT with shaking. Tissue sections were then subsequently incubated overnight with primary antibodies diluted in blocking buffer at 4 °C. Following day, sections were washed three times with 0.2% PBST for 10 minutes each with shaking, and incubated 3hr with secondary antibodies in blocking buffer at RT with shaking. DAPI was added during secondary antibody incubation at a 1:500 dilution. Tissues were then washed with 0.2% PBST three times for 10 minutes each and mounted with Mowiol solution.

### -Immunoblots

The protein lysates from mouse brain tissues were prepared by homogenising the tissues in the RIPA buffer (50mM Tris pH8.0, 150mM NaCl, 1% NP-40, and 0.1% SDS) added with protease (Roche 11836153001) and phosphatase inhibitor cocktails (Roche 04906837001). The protein lysates from cells were directly collected by adding RIPA buffer added with protease and phosphatase inhibitor cocktails. The lysates were mixed with home-made 5x Laemmli sample buffer (5% β-Mercaptoethanol, 0.02% Bromophenol blue, 30% Glycerol 30%, 10% SDS (Sodium dodecyl sulphate), and 250mM Tris pH 6.8) and heated for 5-10 minutes at 95°C. Samples were loaded into the wells in home-made fresh acrylamide gels (6%-13.5%) and directed for SDS–PAGE in Gel-running buffer (33 mM Tris pH 7.4, 192mM Glycine, and 0.2% SDS). Samples were wet-transferred in Transfer buffer (33 mM Tris pH 7.4, 192mM Glycine, and 20% methanol) overnight onto 0.45-μm nitrocellulose membranes (EMD Millipore IPVH00010) at 4 °C. Membranes were incubated 1hr at RT with shaking in blocking solution (5% non-fat skim milk in TBST (20mM Tris pH 8.0, 150mM NaCl, and 0.1% Tween-20)) and incubated overnight with primary antibodies diluted in blocking solution. Membranes were washed 3x with TBST for 10min each with shaking and incubated with secondary antibodies diluted in antibody solution for 1 h at RT. Following 3x wash in TBST, chemiluminescence on membranes acquired using ChemiDoc imaging system (Bio-Rad) running Image Lab software (Bio-Rad). The bands for specific proteins were verified based on the molecular weights aligned with Precision Plus Protein Dual Colour Standards (Bio-Rad 1610374) and analysed using Image Lab software.

### -Immunoprecipitation

For immunoprecipitation, the protein lysates were prepared in IP Lysis buffer (25mM Tris pH 7.5, 150mM NaCl, 1mM EDTA, 0.1% NP-40, 5% Glycerol) added with protease and phosphatase inhibitor cocktails. The lysates were incubated first with antibodies (0.4μg) for an hour and consecutively with Protein A/G PLUS-Agarose beads (SCBT sc-2003) overnight in 1.5ml tubes at 4 °C with rotation. Next day, the lysate mixtures were washed 3x with IP lysis buffer, and the immunoprecipitated lysate samples were subjected to immunoblot using the same procedure described above.

### -Microscope image acquisition and analysis

Leica sp8 confocal microscope was used for image acquisition of neuron cultures, HeLa cells, and most of immunofluorescence imaging on sections. Zeiss Z1 or Zeiss slide scanner Axioscan (10x objective) was used to acquire single plane images of callosal neurons after in utero electroporation (Figure 4). Whole mount cleared tissues were imaged with Zeiss Lightsheet microscope (5x objective). 2D Images were processed, analysed, and measured with NIH ImageJ software. 3D whole mount images were processed, analysed, and measured with Imaris software (Bitplane). For 3d surface generation, machine learning.

### -Statistical analysis

All statistical analyses were performed using GraphPad Prism software. Two groups of measurements were compared by unpaired Student’s t-test. Multiple groups were compared by one-way analysis of variance (ANOVA) followed by Tukey’s or Dunnett’s multiple comparisons test.

### -Ligand binding, clustering, and surface expression/endocytosis assay

For the functional assays, HeLa cell (ATCC CCL-2) was used to achieve sore ectopic EPHB2 receptor expression. Cells were seeded on a coverslip at either 15,000 cells per a well in 24-well plate (Ligand binding and surface expression/endocytosis assay) or seeded at 300,000 cells per a well in 6-well plate (clustering assay) with Dulbecco’s Modified Eagle Medium (DMEM) (Gibco 11965118) supplemented with 10% FBS and 100U/ml Penicillin-Streptomycin (Gibco 15140122) (10% DMEM). Next day, cells were transfected with desired plasmids (1.5μg DNA each for a well in 6well and 0.3ug DNA each for a well in 24 well) in low-serum media Opti-mem (Gibco 31985070) using conventional transfection method (Lipofectamine 3000). Cells were rinsed once with PBS after 6hrs, and media were replaced with 0.5% DMEM. Next day, the cells were treated with pre-clustered recombinant proteins for 30 minutes. Pre-clustering was achieved by mixing 3μg/ml Cy5-conjugated donkey anti-goat IgG (Except clustering assay) and 0.5μg/ml goat anti-human IgG Fc (Millipore Sigma I9135) with either 2μg/ml recombinant human IgG Fc protein (Millipore Sigma 401104) or 2μg/ml recombinant mouse ephrin-B1 Fc chimera protein (R&D Systems 473-EB). After ligand treatment, the rest of the preparation follows the same procedure as described in immunoblots/immunoprecipitation (Clustering assay) or immunocytochemistry (The rest). For surface expression/endocytosis assay, the permeabilization step was skipped.

### -Biolayer interferometry

The receptor-binding domain of human ephrin-B1 (residues 29-166) and human ephrin-B2 (residues 27-167) were cloned into the pHLsec mammalian expression vector (Aricescu et al., 2006) in-frame with a C-terminal hexa-histidine (His6) tag. The complete ectodomain (residues 19-542) and its variant without the ligand-binding domain (ΔLBD) (residues 195-542) were cloned into the pHLsec vector in-frame with a C-terminal 3C-Avi-His6 tag. Protein constructs were produced in HEK 293T (ATCC CRL-3216) cells at 37 °C. The conditioned medium was collected 5 days post transfection and was buffer exchanged using a QuixStand diafiltration system (GE Healthcare). Proteins were purified by immobilised metal affinity chromatography (HisTrap FF column, Cytivia) followed by size-exclusion chromatography (Superdex 200 increase 10/300 GL column, Cytivia). All measurements were performed at 30°C using an Octet Red96e instrument and amine reactive (AR2G) biosensor (both Sartorius). Biosensor surfaces were activated for 6 min with a 1:1 mixture of 10 mM sulfo-NHS (N-hydroxysulfosuccinimide) and 20 mM EDC (3-(N,N-dimethylamino) propyl-N-ethylcarbodiimide). EPHB2 constructs were immobilised as ligands at a concentration of 40 μg/ml in 10 mM sodium acetate, pH 5.0, followed by biosensor blocking in 1 M ethanolamine, pH 8.5. Reference biosensors were immersed into 10 mM sodium acetate, pH 5.0 buffer instead of ligand solution. To collect kinetic binding data, ephrin-B1 or ephrin-B2 analytes were diluted in 10 mM PBS, 1% BSA, 0.01% Tween-20, and 0.05% sodium azide. Multi-cycle kinetic experiments were performed with association and dissociation steps of 300 s and a 1:2 analyte dilution series starting at 800 nM. Data were processed and analysed using the Octet Data Analysis HT software (version 11.1, Sartorius). All kinetic data were double-referenced. Steady-state KD values were determined from fitting a Langmuir 1:1 binding isotherm to the maximum response unit values in OriginPro 9.1. Experiments and analyses were performed in duplicates.

### -*In utero* electroporation

E15.5 CD1 or B6D2F1 pregnant females were used for in utero electroporation. The female mice were introduced with 0.05ml buprenorphine by subcutaneous injection several hours prior to the surgery. The mice were laid on a warm pad and anesthetised by isoflurane (CDMV 108737) inhalation during the surgery. The abdominal skin and wall were cut, and the uterus containing embryos was pulled out. Then, 1μl of DNA solution including plasmids (up to 3μg; 1μg/1μl for vector, shRNA, and OE plasmids; 0.5μg/1μl for GFP or RFP expressing plasmids) and 0.03% Fast Green was injected with a glass micropipette (Drummond Scientific 5-000-2005) into the left lateral ventricle of the embryos. The head of the embryos was placed between 5mm tweezer electrodes (NepaGene CUY650P5) at about 90° angle to target somatosensory cortex (Jan Baumgart and Nadine Baumgart, Jove, 2016), and two poring electric pulses (40V, 50ms pulse length, 950ms pulse interval, and 10% decay rate) and three transfer electric pulses (35V, 50ms pulse length, 950ms pulse interval, and 0% decay rate) were sequentially delivered using NEPA21 electroporator (NepaGene). After the electroporation, the uterus was returned into the abdominal cavity. Abdominal wall was sutured, and the skin was closed with 9mm Autoclips (Braintree Scientific ACS CS 0009952). During the entire surgery, the mice were applied with warm light to maintain the body temperature within 36-37 °C and consistently moisturised with warm PBS around the incision site and uterus to prevent dehydration. After surgery, the mice were kept in recovery chamber with warm light illumination for 15-30m and returned to the cage.

### -Primary neuron culture

E16.5 CD1 or B6J mouse embryos were used for all primary cortical neuron cultures. Cortices were dissected in CMF-BSS buffer (500ml Distilled Water (Gibco 15230162), HBSS (Gibco 14185052), HEPES (Gibco 15630080), and Penicillin-Streptomycin (Gibco 15140122)), followed by digestion with 0.25% trypsin (Gibco 25200072) for 30 min at 37 °C. Tissues were washed more than three times with CMF-BSS buffer and dissociated in DMEM with 10% fetal bovine serum (FBS) (Gibco 12483020) by gently pipetting up and down 10 times with a Pasteur pipette (VWR 14672380). Dissociated cells were filtered through a 70μm cell strainer (Fisher Scientific 08-771-2) and plated onto 1mg/ml Poly-L-Lysine-coated (Millipore Sigma P2636) 1.5-mm coverslips (Fisher Scientific NC0308920) in 12-well plates for Immunostaining or 6-well plates (Sarstedt 83.3920) for biochemistry. The dissociated cells were first cultured in plating medium (500ml MEM (Millipore Sigma M4655 or Gibco 11095-080), glucose, pyruvate, and horse serum (Gibco 26050070)). 3h later, the medium was replaced with Neurobasal medium (Gibco 21103049) supplemented with SM1 Neuronal Supplement (Stemcell 05711), and GlutaMAX-I (Gibco 35050). For immunostaining, neurons were plated at 100,000 cells (500,000 after nucleofection) per a well in 12-well plate and cultured for various periods up to 7d. For biochemistry, neurons were plated at 2M (5M after nucleofection) per a well in 6-well plate and cultured for 6d with 500μl addition of Neurobasal at DIV3. Drug treatments were carried out 24hr prior to the fixation at indicated concentrations.

### -Nucleofection

To achieve an optimal transfection of dissociated neurons, commercial nucleofection kit (Lonza) was used. After dissociating primary neurons, desired number of neurons (5M per sample) were transferred to a 15ml conical tube and centrifuged at 1,500 rpm for 5 minutes. After removing supernatant, the cells were resuspended with Lonza buffer (100μl/sample) and mixed with plasmids (8μg in total per sample). The mixture (100μl) was transferred to a nucleofection cuvette and received electroporation using a Lonza Nucleofector (Protocol: O-05). 950μl plating medium was directly added into a cuvette, and the transfected neurons were plated at a desired density.

### -iDISCO+ wholemount tissue clearing and corpus callosum volumetric analysis

An optimised iDISCO+ protocol was used for tissue clearing of young and adult mouse brains^87^. All brain tissues for tissue clearing were post-fixed overnight with 4% PFA and washed 3x with PBS. The entire brain was used for tissue clearing of P4 mice, and the forebrain tissues were used for P14 and older mice. The brain tissues were first dehydrated by incubating in sequential gradient of 20%, 40%, 60%, 80%, 100%, and 100% methanol/water solution for 1hr each at RT. The tissues were then incubated in de-lipidation solution (66% dichloromethane and 33% methanol) overnight at RT with shaking. The tissues were washed 2x with 100% methanol for 1hr each with shaking and incubated with bleaching solution (5% H2O2 in methanol) overnight at 4 °C. Next day, the samples were rehydrated with sequential gradient of 80%, 60%, 40%, 20% methanol/water solution, and twice with PBS for 1hr each at RT. The tissues were sequentially permeabilized in PBSGT (PBS with 0.2% gelatine and 2% triton x-100) added with 0.3M glycine and blocked in PBSGT added with 5% IHS for 2 and 3 days respectively at 37°C with rotation. After blocking tissues were cut leaving desired regions. The tissues were incubated with primary antibodies diluted in PBSGT added with 1% IHS and 0.1% saponin for 1 week (P4) or 3 weeks (>P13) at 37°C with rotation. Tissues were washed 6x with PBS for 1h each at RT with shaking and incubated with secondary antibodies in PBSGT added with 1% IHS and 0.1% saponin for 1 week at 37°C with rotation. After 6x wash with PBS for 1h each, the tissues were again sequentially dehydrated and incubated with de-lipidation solution for 3hr at RT with shaking. Tissues were washed 2x with dichloromethane for 15min each at RT with shaking and incubated in dibenzyl ether until imaging. During the entire process, the tissues were maintained without (minimum) light exposure.

### -Single-cell RNA sequencing data analysis

To identify CPNs in the developing mouse brain atlas of La Manno et al.^53^ (E7–E18; “pooled” counts normalised to the median library size), we first selected individual cells and then the atlas clusters to which they belonged. Candidate cells were dorsal forebrain neurons sampled between E15.0 and E18.0 (17,917 cells, 131 clusters). Cells were characterised with the CellCover marker gene panels^88^, derived from cortical cells labelled 1, 24 and 96 hours after their last division at E12–E15^89^; the E15.96H panel marks young superficial-layer neurons. In each cell, one-sided Wilcoxon rank-sum tests compared E15.96H panel genes with E15.24H panel genes and with deep-layer markers (Tbr1, Bcl11b, Fezf2). Retained cells expressed Satb2, Pou3f2 or Trim67, were not cycling, had a higher E15.96H than E15.24H signal (Bonferroni-corrected P < 0.05) and deep-layer signal (P < 0.05, or no deep-layer marker), and belonged to the cortical or hippocampal glutamatergic subclass (3,004 cells, 30 clusters). Six of these clusters (Neur512–Neur517), with high Satb2, low Fezf2 and Bcl11b and the E15.96H signature, were taken as CPN clusters, and all their cells (6,682) were used in Fig. 4a and Extended Data Fig. 4a (mean ± SEM/age). The workflow to reproduce these results from the public atlas is available at https://github.com/maunei/callosal_projection_neurons_selection.

### -Mutant mouse modelling

*Ephb2* C97R and *Ephb2* K873N knock-in mouse lines were generated using CRISPR-Cas9 system via i-GONAD procedure^90^. Briefly, recombinant Cas9 nuclease (IDT cat# 1081060) and guide RNAs (gRNAs), consisting of CRISPR RNAs (crRNAs) and trans-activating CRISPR RNAs (tracrRNAs) (IDT cat# 1072532), were preassembled into ribonucleoprotein (RNP) complexes and co-injected with the corresponding single-stranded oligodeoxynucleotide (ssODN) donor templates containing the desired mutation into the ampulla of CD1 pregnant females at 0.7 days post coitum, followed by electroporation. Mice were genotyped by PCR followed by Sanger sequencing to confirm the presence of the intended mutations. The CRISPR-Cas9 gRNA design tool from IDT were used to select gRNAs. All sequences used are listed in Supplementary Table 3. Before experimental use, all mutant mice were backcrossed with C57BL/6J (B6J) mice for more than six generations.

## Acknowledgements

We thank all the families who participated in this study for sharing clinical data and allowing data publication. We thank M. Greenberg, S. Margolis, K. Zito, P. Penzes, E. Parnell for sharing experimental reagents including antibodies and plasmids. We thank M. Liang for technical assistance. We thank IRCM Core Facility employees: O. Jumanca, M. Laprise, and animal facility technicians for animal cares; L. Lian from the Disease Modelling and Genome Editing platform for expert technical assistance in the generation of the mouse models; D. Filion and M. Duguay for microscopy assistance supported, in part, by funding from the Canadian Foundation for Innovation. S.S. Park received scholarships and travel awards from Fondation IRCM and McGill University. A. Kania was supported by Canadian Institutes for Health Research supported with Project and Operating Grants PJT-162225, MOP-77556, PJT-153053, PJT-159839, and PJT-197987, the Doggone Foundation Chair of Excellence in Pain and the Fondation IRCM. G.N. Nagy was supported by the János Bolyai Research Fellowship of the Hungarian Academy of Sciences and the Mihály Somogyi Grant of the Varga József Foundation. E.Y. Jones and G.N. Nagy were supported by the Medical Research Council UK (MR/T000503/1) and the Sir Andrew McMichael Chair of Structural Immunology. Work in the laboratory of E. Herrera is supported by the Spanish State Research Agency (PID2025-168114NB-I00), the PROMETEO Program of the Generalitat Valenciana (CIPROM/2023/15), and the “Severo Ochoa” Program for Centres of Excellence in R&D (CEX2021-001165-S funded by MICIU/AEI/ 10.13039/501100011033). I. Pérez Ferrer holds a fellowship from the Spanish Ministry of Science, Innovation and Universities (PRE2021-098605). Exome sequencing in proband 1 was performed as part of the “neuromendeliome” research project financially supported by Agence de Biomedecine. L. Caporali was supported by grants GR-2016-02361449 from the Italian Ministry of Health. D. Bauer participated in initial modelling of C97R variant from proband 1.

## Author contributions

S.S.P. and A.K. designed the study. G.N.N. performed structure analysis with input from E.Y.J. S.S.P., G.N.N., S.K., and B.Z. performed cell function assays and the data analyses. L.L. and P.O. performed human genetic analyses. M.N. and P.O. performed bioinformatics. S.S.P. and C.K. performed mouse brain anatomy, immunohistochemistry, immunocytochemistry, biochemistry, imaging, and data analyses. I.P.F., Y.C., M.H., S.S.P. performed in utero electroporation, imaging, and data analyses with input from E.H. Y.K. generated CRIPSR/Cas9-mediated knock-in mice and S.S.P. performed mouse breeding, immunohistochemistry, and data analyses. C.D., P.O., F.P., V.C., G.D.H, K.A., M.W.E., and their teams generated and analysed the exome sequencing data and identified the variants in patients. S.S.P., G.N.N., and A.K. wrote the manuscript.

## Additional information

The authors declare no competing financial or other interests.

---

**Extended Data Fig. 1:**
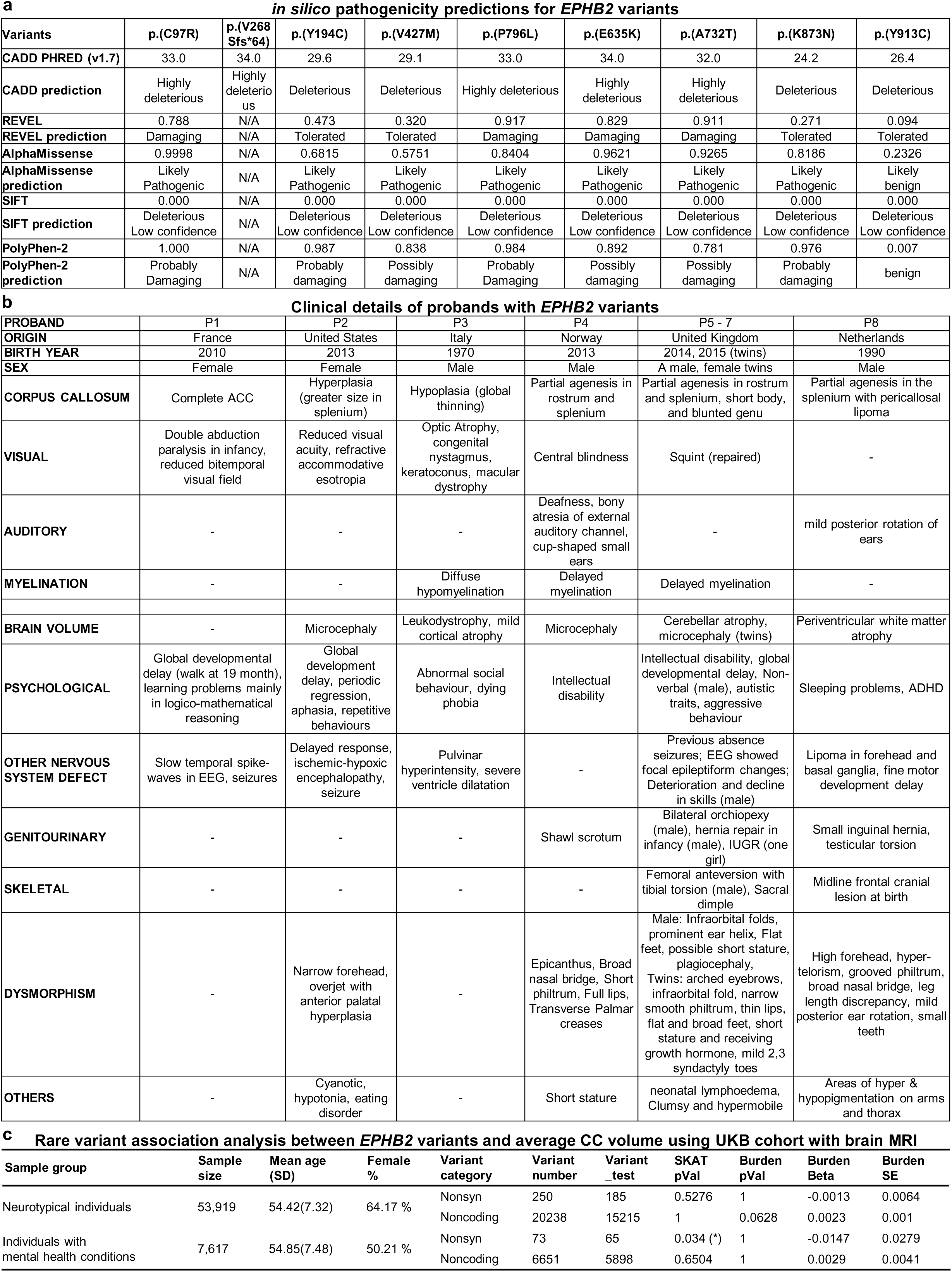
Genetic and clinical information on *EPHB2* variants and the carriers. **a**, Table summarising *in silico* pathogenicity predictions and functional annotation profiles of *EPHB2* variants. Pathogenicity predictions and qualitative classifications are based on the established cut-off criteria of each respective bioinformatic tool. CADD: Combined Annotation Dependent Depletion (v1.7), REVEL: Rare Exome Variant Ensemble Learner, AlphaMissense: Google DeepMind AlphaMissense (v1.0), SIFT: Sorting Intolerant From Tolerant, PolyPhen-2: Polymorphism Phenotyping v2, N/A: Not Applicable (due to the variant type being a frameshift/truncating mutation). b, Table summarizing de-identified clinical information of human probands carrying *EPHB2* variants. **b,** Table summarising clinical information based on case reports from individuals with *EPHB2* variants. **c,** Table summarising the rare variant association analysis between EPHB2 variants and average CC volume using UKB cohort with brain MRI. Inclusion and exclusion criteria are described in the method. The significance was adjusted with Bonferroni correction. * P<0.05

**Extended Data Fig. 2:**
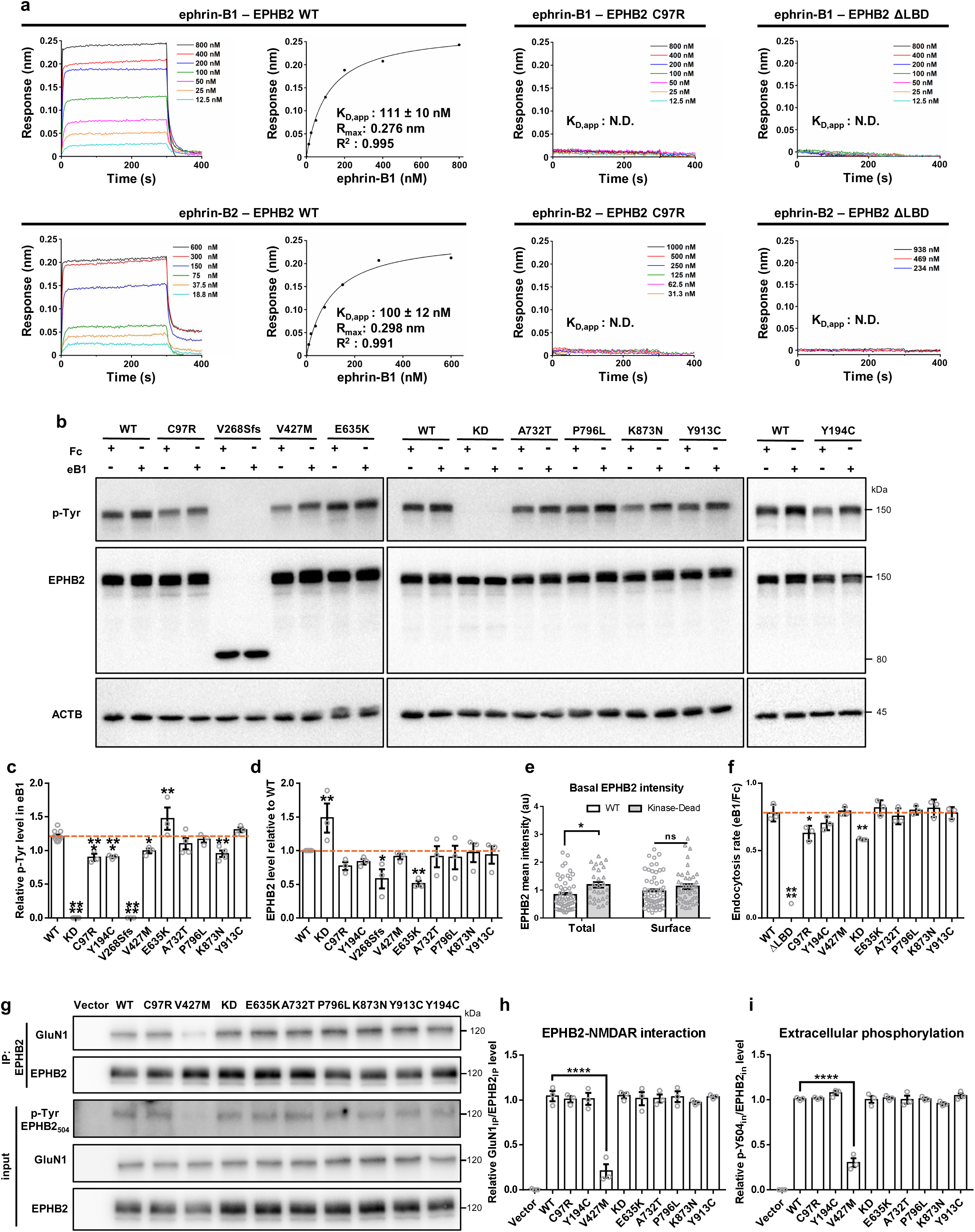
EPHB2 variants impair distinct EPH signalling steps. **a**, Kinetics of EPHB2 ectodomain interaction with RBD domain of ephrin-B1 or ephrin-B2 using Biolayer Interferometry (means ± SEM, n = 2). **b,** Immunoblot images of HeLa cell lysates transfected with EPHB2 variants and treated with either Fc IgG fragment or pre-clustered ephrin-B1 for 30 minutes. **c,d,** Quantification of relative phospho-tyrosine (c) and EPHB2 (d) levels in HeLa cells transfected with EPHB2 variants. SEM; N>4. **e,** Quantification of total WT-EPHB2 and KD-EPHB2 protein expressed in HeLa cells, detected via anti-GFP and anti-EphB2 ectodomain antibodies in non-permeabilising conditions, expressed as mean signal intensity. Each data point represents an individual cell. N>50 from three independent experiments. **f,** Quantification of the endocytosis rate measured as the surface/total EPHB2 level treated with pre-clustered ephrin-B1 normalised to the surface/total EPHB2 level treated with Fc IgG fragment. **g,** Immunoblots of GluN1 co-immunoprecipitation with EPHB2 and input lysates from HEK293T cells transfected with EPHB2 variants. **h,** Quantification of relative GluN1 level normalised to EPHB2 level in IP lysates (GluN1_IP_/EPHB2_IP_) from HEK293T cells transfected with EPHB2 variants. **i,** Quantification of relative EPHB2 extracellular phospho-tyrosine (p-Y504) signal normalised to EPHB2 levels in input lysates (p-Y504_in_/EPHB2_in_) from HEK293T cells transfected with EPHB2 variants.

**Extended Data Fig. 3:**
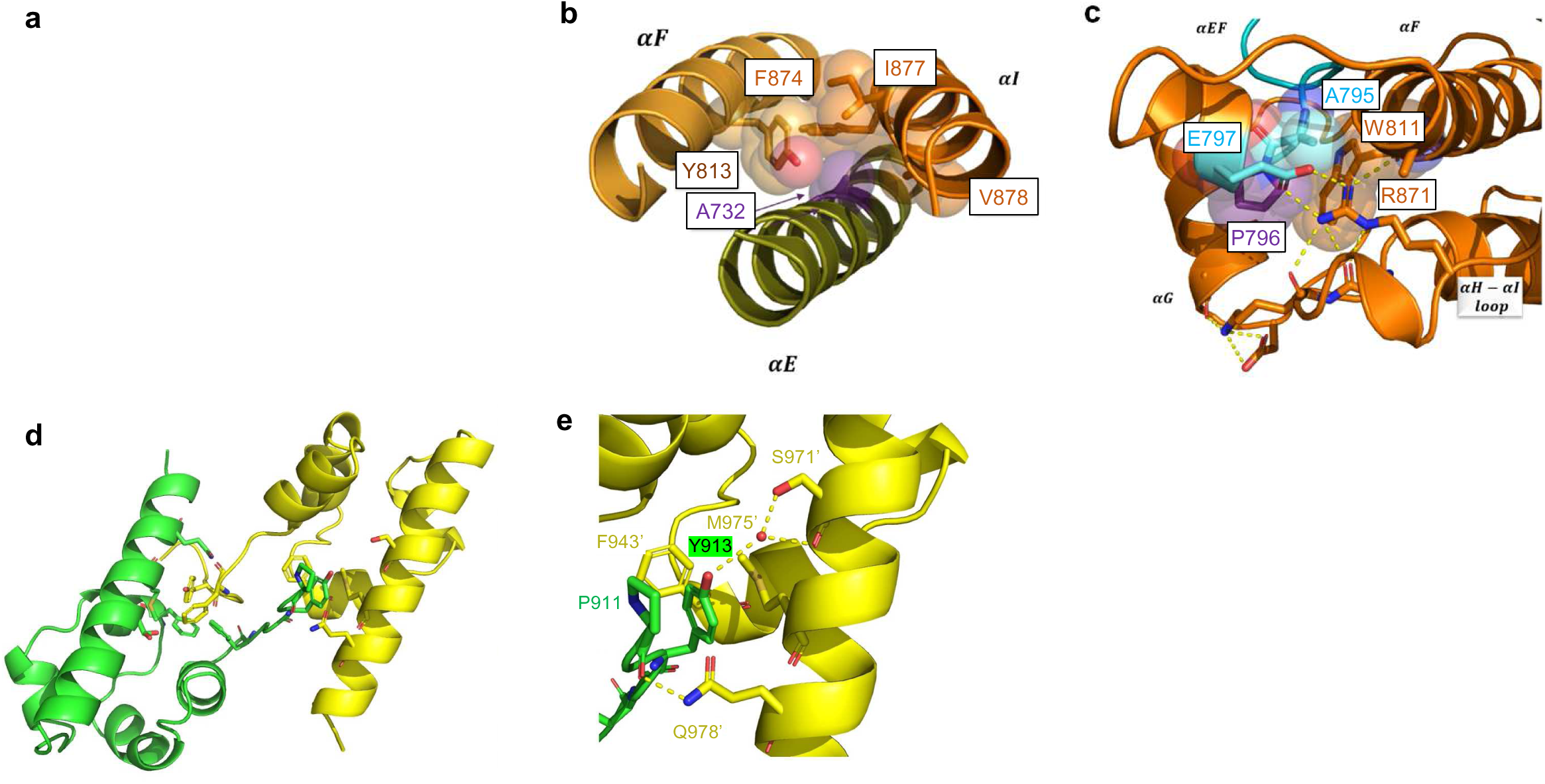
Structural analysis of additional EPHB2 variants. **a, b,** A732 (purple) in helix αE from the kinase C-lobe, packing against Y813 and F874 (αF) and I876 and V877 (αI). **c,** P796 (purple) at the C-terminal end of the activation loop (teal) in the 5I9W-based model, contacting A794, W810, E796 and R870; hydrogen bonds are dashed lines**. d,** EPHB2 SAM domain arm-exchange dimer (PDB 1B4F; chains green and yellow). **e,** Close-up of Y913 buried in a cleft formed by P911 of the same chain and F943′, S971′, M975′ and Q978′ of the partner chain (primed labels).

**Extended Data Fig. 4:**
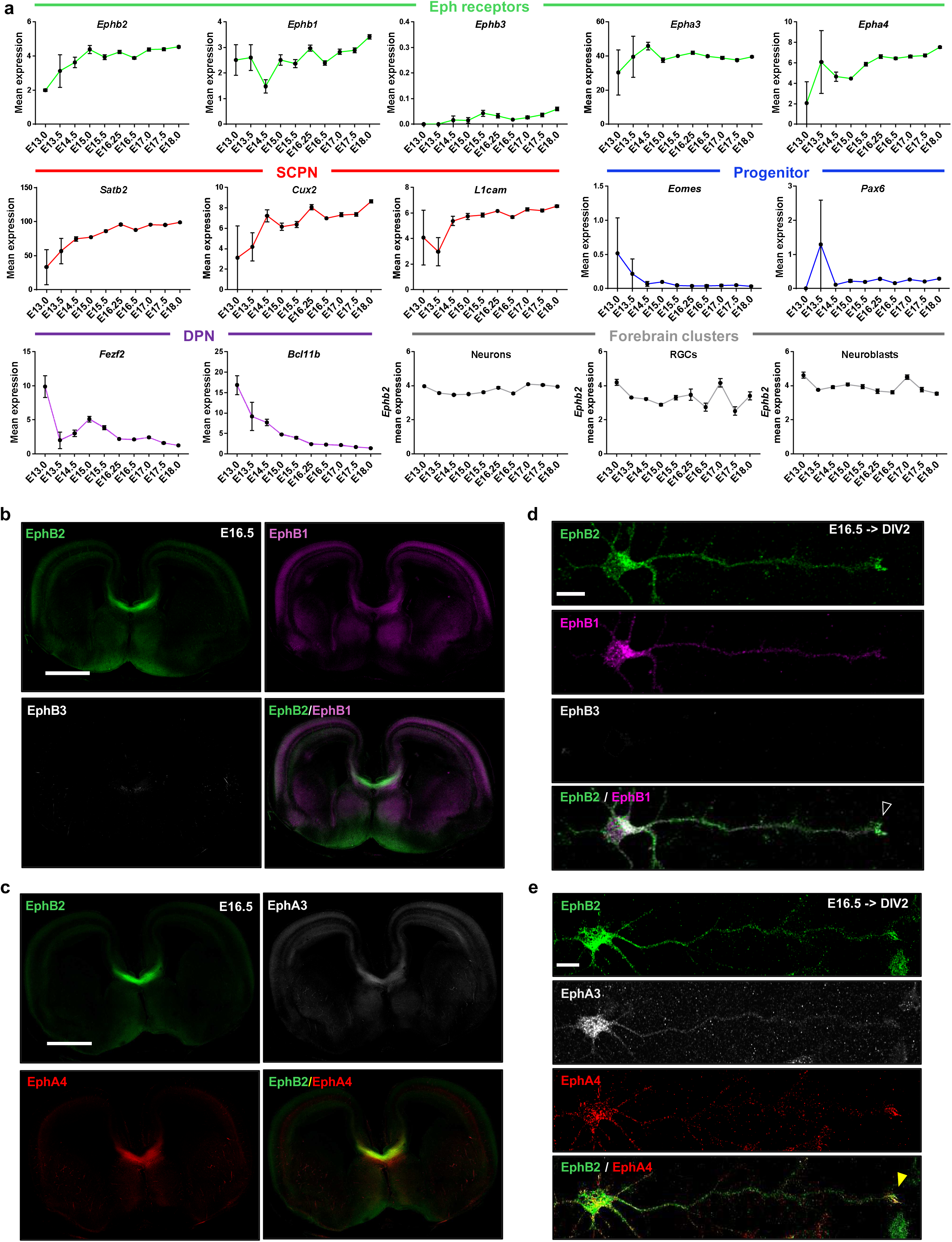
Embryonic expression of Eph receptors, CPN and progenitor markers. **a**, Mean expression of genes within CPN clusters and Ephb2 expression within pooled forebrain clusters from E13-E18 mouse forebrains. The data were generated using single-cell RNA-seq datasets from PMID 34321664 (La Manno et al., Nature, 2021). SCPN indicates superficial callosal projection neuron markers. DPN indicates deep layer projection neuron markers. The CPN clusters include 6,682 cells in total. SEM; N=2 (E13.0), 5 (E13.5), 70 (E14.5), 188 (E15.0), 372 (E15.5), 717 (E16.25), 1,318 (E16.5), 1,041 (E17.0), 1,035 (E17.5), 1,934 (E18.0). Pooled forebrain neuron clusters include 41,552 cells in total (SEM; N=1,481 (E13.0), 2,607 (E13.5), 4,433 (E14.5), 4,805 (E15.0), 2,966 (E15.5), 2,770 (E16.25), 5,209 (E16.5), 6,077 (E17.0), 3,267 (E17.5), 7,937 (E18.0)). Pooled forebrain RGC clusters include 5,001 cells in total (SEM; N=212 (E13.0), 1,970 (E13.5), 1,266 (E14.5), 852 (E15.0), 266 (E15.5), 54 (E16.25), 82 (E16.5), 130 (E17.0), 60 (E17.5), 109 (E18.0)). Pooled forebrain neuroblast clusters include 7,219 cells in total (SEM; N=356 (E13.0), 1,130 (E13.5), 1,166 (E14.5), 915 (E15.0), 452 (E15.5), 393 (E16.25), 705 (E16.5), 645 (E17.0), 431 (E17.5), 1026 (E18.0)). **b,c,** E16.5 mouse forebrain sections stained for EphB2 and EphB1/EphB3 (b) or EphA3/EphA4 (c). Scale bar, 1 mm. **d,e,** Primary cortical neurons at DIV2 stained for EphB2 and EphB1/EphB3 (d) or EphA3/EphA4 (e). Arrowheads indicate the axonal tips. Scale bar, 10 µm.

**Extended Data Fig. 5:**
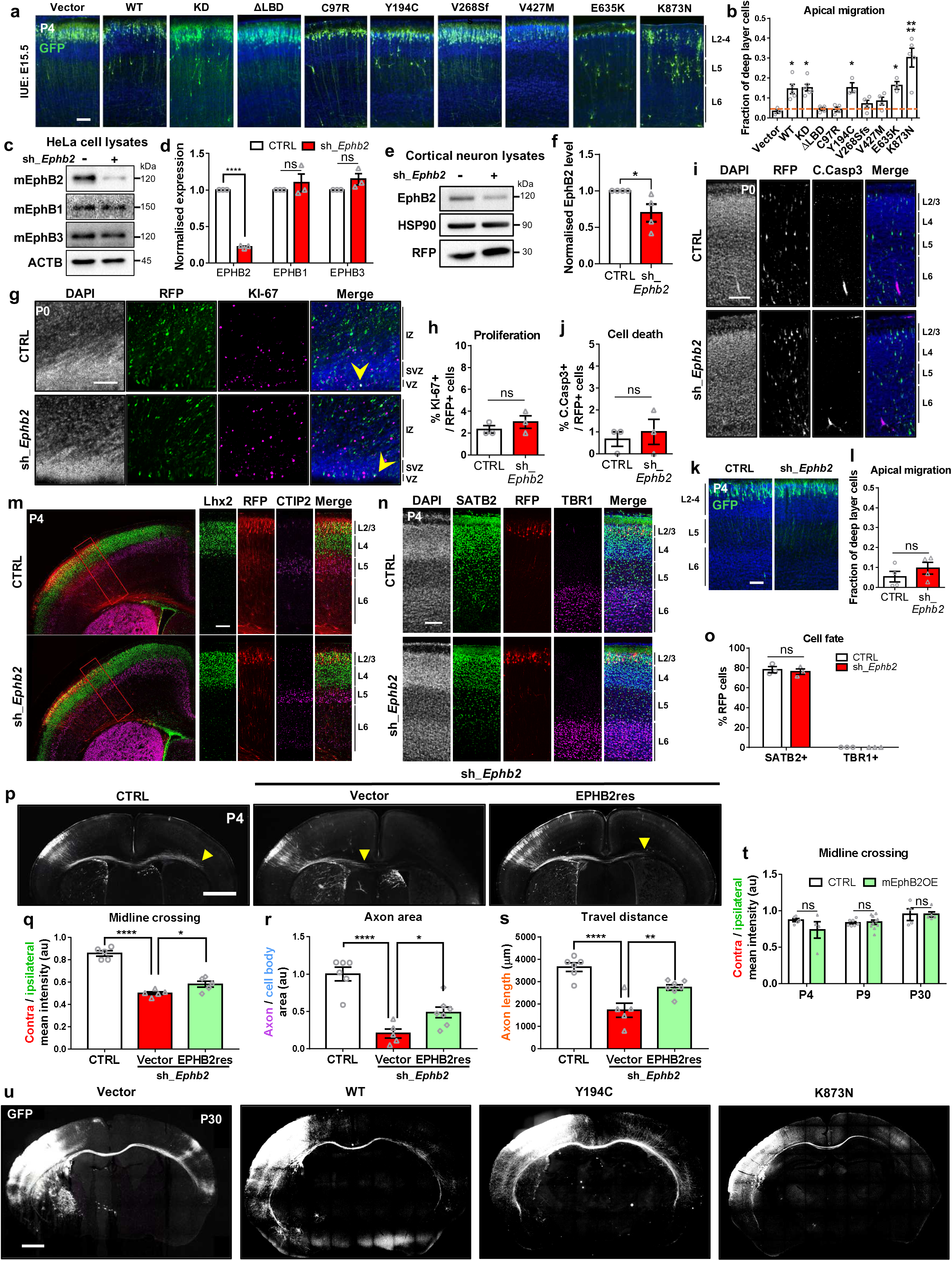
Validation of EphB2 depletion and characterisation of IUE experiments. **a**, Cortical cells co-expressing EPHB2 variants (or vector controls) and GFP in coronal sections of P4 brains. Scale bar, 100 µm. **b,** Quantification of the fraction of GFP cells fund in deep layers. **c,** Detection of EphB proteins and actin (ACTB) in lysates of HeLa cells transfected with plasmids encoding mouse EphB1, EphB2, and EphB3, co-transfected with control plasmids or the sh_*Ephb2* plasmid. **d,** Quantification of Eph receptor levels in c, normalised to actin (ACTB) levels. SEM; N=3. **e,** Detection of EphB2, RFP and HSP90 in DIV6 mouse primary cortical neuron lysates following nucleofection with RFP expression plasmids, and control plasmids or the sh_*Ephb2* plasmid. **f,** Quantification of EphB2 expression e, normalised to HSP90. SEM; N=4. g-o, Examination of proliferation, cell death and cell fate in brains of mice co-electroporated with an RFP or GFP expression plasmid, and control or the sh_*Ephb2* plasmid. **g,** KI-67-expression in proliferating cells and RFP expression in P0 mouse brain sections. Scale bar, 100 µm. Arrowheads indicate RFP and KI-67 co-expressing cells. **h,** Fraction of RFP-expressing cells that also express KI-67 at P0. SEM; N=3 (each replicate includes >50 cells). **i,** Cleaved caspase-3 expression (cell death marker) in RFP-expressing cells in P0 coronal section of the cortex. Scale bar,100 µm. **j,** Fraction of RFP-expressing cells that co-expressed cleaved caspase-3 at P0. SEM; N=3 (each replicate includes >100 cells). **k,** Location of GFP-expressing cells co-electroporated with control or sh_*Ephb2* plasmids in the P4 cortex. Scale bar, 100 µm. **l,** Fraction of GFP-expressing cells found in deep cortical layers at P4. **m,** Lhx2 (L2/3 marker), Ctip2 (L6 marker), and RFP expression in the P4 mouse cortex. Scale bar,100 µm. **n,** Satb2 (callosal marker), Tbr1 (L6 marker), and RFP expression in the P4 mouse cortex. Scale bar,100 µm. **o,** Quantification of the fraction of Satb2- or Tbr1-expressing electroporated (RFP-expressing) cells. SEM; N=3 (each replicate includes >50 cells). **p,** Coronal sections of P4 brains of mice following IUE of sh_*Ephb2* controls plasmid, or the sh_*Ephb2* plasmid and empty vector or a plasmid expressing a sh_*Ephb2*-resistant EPHB2 (EPHB2res). Arrowheads indicate the terminus of the longest axons. **q-s,** Quantification of midline crossing rate (q), normalised axon volume (r), and axon travel distance (s) in the shRNA rescue experiment in p. SEM; N=5-7. **t,** Quantification of midline crossing rate in P4, P9, and P30 brain sections following CTRL or mEphB2 IUE. SEM; N = 4-5 (P4), 7-10 (P9), 4-5 (P30). **u,** Images showing P30 brain sections following IUE with control plasmids and those encoding WT-EPHB2, Y194C-EPHB2, and K873N-EPHB2.

**Extended Data Fig. 6:**
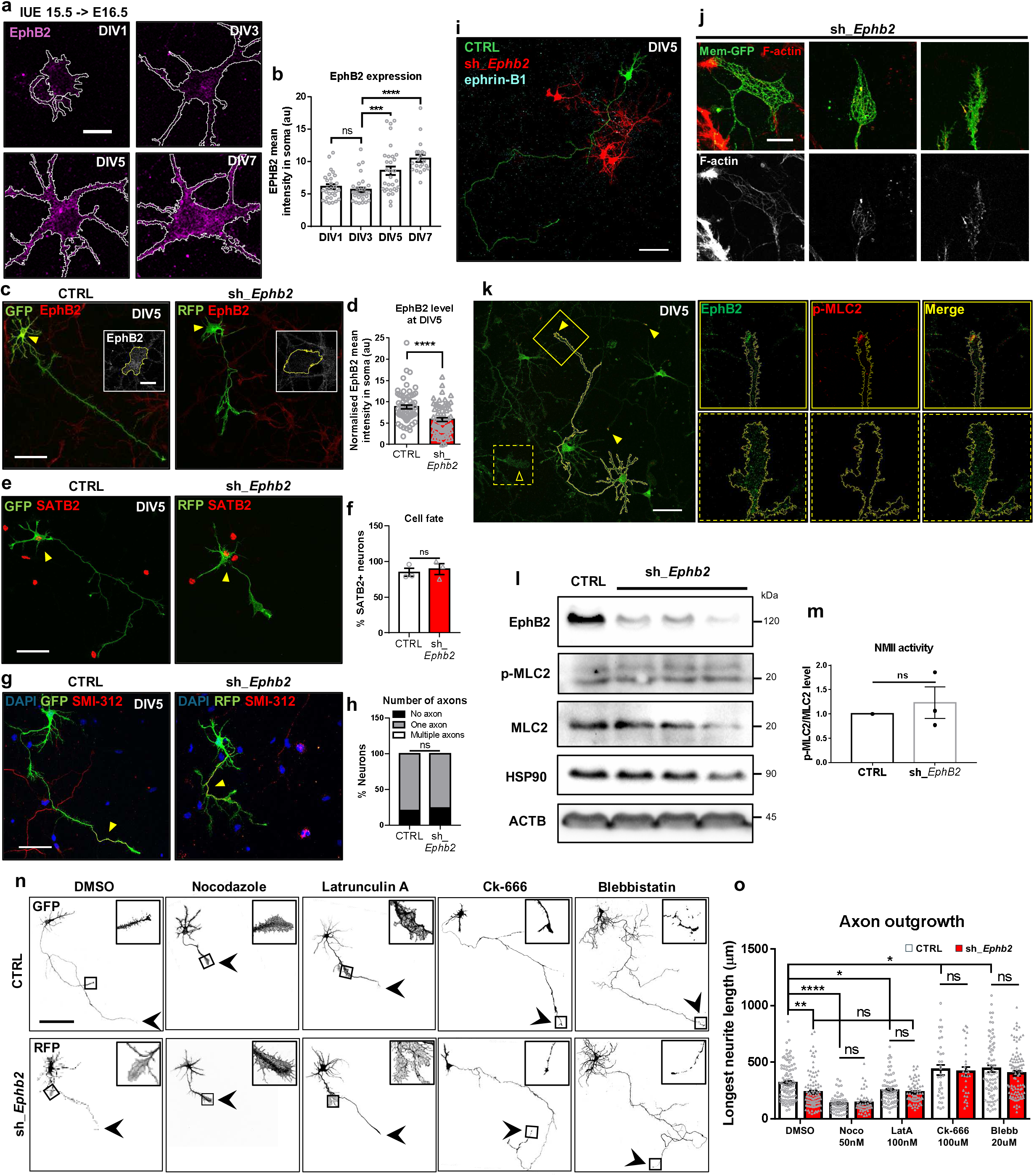
EphB2 promotes callosal axon outgrowth through spatial restriction of actin-myosin at growth cone. **a**, EphB2 localisation in primary neurons electroporated with the control shRNA plasmid at DIV1-7. Scale bar, 10 µm. **b,** Quantification of normalised EphB2 signal intensity in cell bodies (arrowheads) of electroporated primary neurons in a. SEM; N>30 neurons from 3 independent experiments. **c,** EphB2 localisation in DIV5 primary neurons electroporated with either control or sh_*Ephb2* plasmids. Boxed inset shows the cell body outlined in yellow. Scale bar, 50 µm, and 10 µm in inset. **d,** Quantification of normalised EphB2 mean intensity in cell bodies of electroporated primary neurons. SEM; N>100 neurons from 3 independent experiments. **e,** Satb2 expression (arrowheads) in DIV5 primary neurons electroporated with either control or sh_*Ephb2* plasmids. Scale bar, 50 µm. **f,** Quantification of the fraction of electroporated neurons expressing Satb2. SEM; N>50 neurons from 3 independent experiments. **g,** Localisation of the axon specific neurofilament SMI-312 (arrowheads) in DIV5 primary neurons electroporated with either control or sh_*Ephb2* plasmids. Scale bar, 50 µm. **h,** Quantification of the proportion of neurons with the indicated number of axons. SEM; N>50 neurons from 3 independent experiments. **i,** Images of DIV5 primary neurons cultured on ephrin-B1-coated coverslips following electroporation with either control or sh_*Ephb2* plasmids. Scale bar, 50 µm. **j,** Localisation of membrane linked GFP (mem-GFP) and F-actin in axonal termini of DIV5 primary neurons electroporated with the sh_*Ephb2* plasmid. Scale bar, 10 µm. **k,** Localisation of EphB2 and p-MLC2 in DIV5 nascent primary neurons. Filled arrowheads indicate EphB2 and p-MLC2 co-localisation. Scale bar, 50 µm. **l,** Immunoblots of cell lysates from DIV6 rat primary cortical neurons following nucleofection with either with control or sh_*Ephb2* plasmids. **m,** Quantification of the ratio of p-MLC2 to MLC2 from rat primary cortical neuron lysates following nucleofection with either control or sh_*Ephb2* plasmids. **n,** Images of DIV5 primary neurons electroporated with either control or sh_*Ephb2* plasmids following 1 day of treatment with the indicated drugs. Scale bar, 100 µm. **o,** Quantification of the longest neurite length in DIV5 primary neurons electroporated with either control or sh_*Ephb2* plasmids following drug treatment. SEM; N=30-100 neurons from 3 independent experiment.

**Supplementary Table 1:** Full list of *EPHB2* rare variants in individuals with mental health conditions in UKB cohort.

| no. | DNA (GRCh38) | Isoform | Exon | protein | domain | Allele frequency | No. Carrier |
|---|---|---|---|---|---|---|---|
| 1 | chr1:22784429:T:C | NM_004442.7 | 3/16 | p.T55M | LBD | 5.005E-05 | 1 |
| 2 | chr1:22784454:A:C | NM_004442.7 | 3/16 | p.V63K | LBD | 4.997E-05 | 1 |
| 3 | chr1:22784455:A:G | NM_004442.7 | 3/16 | p.V64M | LBD | 5.068E-05 | 1 |
| 4 | chr1:22784504:A:G | NM_004442.7 | 3/16 | p.R80H | LBD | 9.965E-05 | 2 |
| 5 | chr1:22784506:T:C | NM_004442.7 | 3/16 | p.R81C | LBD | 4.983E-05 | 1 |
| 6 | chr1:22784521:G:A | NM_004442.7 | 3/16 | p.I86V | LBD | 4.974E-05 | 1 |
| 7 | chr1:22784549:A:G | NM_004442.7 | 3/16 | p.R95H | LBD | 4.981E-05 | 1 |
| 8 | chr1:22784651:G:A | NM_004442.7 | 3/16 | p.N129S | LBD | 4.979E-05 | 1 |
| 9 | chr1:22784728:T:C | NM_004442.7 | 3/16 | p.R155C | LBD | 4.967E-05 | 1 |
| 10 | chr1:22784746:G:A | NM_004442.7 | 3/16 | p.T161A | LBD | 4.961E-05 | 1 |
| 11 | chr1:22784747:G:C | NM_004442.7 | 3/16 | p.T161S | LBD | 4.960E-05 | 1 |
| 12 | chr1:22784749:A:G | NM_004442.7 | 3/16 | p.E162K | LBD | 4.960E-05 | 1 |
| 13 | chr1:22784767:T:C | NM_004442.7 | 3/16 | p.P168S | LBD | 9.945E-05 | 2 |
| 14 | chr1:22784812:A:G | NM_004442.7 | 3/16 | p.G183S | LBD | 4.955E-05 | 1 |
| 15 | chr1:22784833:A:G | NM_004442.7 | 3/16 | p.V190M | LBD | 4.951E-05 | 1 |
| 16 | chr1:22784933:A:G | NM_004442.7 | 3/16 | p.R223Q | CRD | 4.951E-05 | 1 |
| 17 | chr1:22784963:A:T | NM_004442.7 | 3/16 | p.V233E | CRD | 4.950E-05 | 1 |
| 18 | chr1:22785016:G:A | NM_004442.7 | 3/16 | p.I251V | CRD | 4.958E-05 | 1 |
| 19 | chr1:22785052:A:G | NM_004442.7 | 3/16 | p.V263I | CRD | 4.974E-05 | 1 |
| 20 | chr1:22785074:A:G | NM_004442.7 | 3/16 | p.R270Q | CRD | 4.980E-05 | 1 |
| 21 | chr1:22863072:C:G | NM_004442.7 | 4/16 | p.D283H | CRD | 3.016E-04 | 6 |
| 22 | chr1:22863090:C:T | NM_004442.7 | 4/16 | p.C289R | CRD | 5.043E-05 | 1 |
| 23 | chr1:22864951:T:C | NM_004442.7 | 5/16 | p.R348C | FNIII-1 | 5.095E-05 | 1 |
| 24 | chr1:22864954:A:G | NM_004442.7 | 5/16 | p.D349N | FNIII-1 | 5.095E-05 | 1 |
| 25 | chr1:22864990:G:A | NM_004442.7 | 5/16 | p.I361V | FNIII-1 | 1.106E-03 | 22 |
| 26 | chr1:22865009:T:C | NM_004442.7 | 5/16 | p.S367L | FNIII-1 | 1.014E-04 | 2 |
| 27 | chr1:22865014:T:C | NM_004442.7 | 5/16 | p.R369W | FNIII-1 | 1.014E-04 | 2 |
| 28 | chr1:22865015:A:G | NM_004442.7 | 5/16 | p.R369Q | FNIII-1 | 6.082E-04 | 12 |
| 29 | chr1:22865026:G:A | NM_004442.7 | 5/16 | p.T373A | FNIII-1 | 5.009E-05 | 1 |
| 30 | chr1:22865030:A:G | NM_004442.7 | 5/16 | p.R374H | FNIII-1 | 5.009E-05 | 1 |
| 31 | chr1:22865053:A:G | NM_004442.7 | 5/16 | p.A382T | FNIII-1 | 3.997E-04 | 8 |
| 32 | chr1:22865080:G:C | NM_004442.7 | 5/16 | p.P391A | FNIII-1 | 5.030E-05 | 1 |
| 33 | chr1:22865131:G:A | NM_004442.7 | 5/16 | p.I408V | FNIII-1 | 1.009E-04 | 2 |
| 34 | chr1:22882394:T:C | NM_004442.7 | 6/16 | p.R447C | FNIII-2 | 4.980E-05 | 1 |
| 35 | chr1:22882400:A:G | NM_004442.7 | 6/16 | p.V449M | FNIII-2 | 4.980E-05 | 1 |
| 36 | chr1:22882448:A:G | NM_004442.7 | 6/16 | p.V465M | FNIII-2 | 9.973E-05 | 2 |
| 37 | chr1:22882470:C:A | NM_004442.7 | 6/16 | p.Q472P | FNIII-2 | 4.957E-05 | 1 |
| 38 | chr1:22882478:A:G | NM_004442.7 | 6/16 | p.E475K | FNIII-2 | 4.957E-05 | 1 |
| 39 | chr1:22892938:A:G | NM_004442.7 | 7/16 | p.V495M | FNIII-2 | 2.520E-04 | 5 |
| 40 | chr1:22892956:A:G | NM_004442.7 | 7/16 | p.G501S | FNIII-2 | 1.519E-04 | 3 |
| 41 | chr1:22892980:T:C | NM_004442.7 | 7/16 | p.R509W | FNIII-2 | 5.028E-05 | 1 |
| 42 | chr1:22895523:T:C | NM_004442.7 | 8/16 | p.S548L | TM | 4.984E-05 | 1 |
| 43 | chr1:22895537:A:G | NM_004442.7 | 8/16 | p.V553I | TM | 4.972E-05 | 1 |
| 44 | chr1:22896415:T:C | NM_004442.7 | 9/16 | p.R569W | JM | 3.512E-04 | 7 |
| 45 | chr1:22896427:T:C | NM_004442.7 | 9/16 | p.R573C | JM | 9.997E-05 | 2 |
| 46 | chr1:22905992:G:C | NM_004442.7 | 10/16 | p.P592A | JM | 5.032E-05 | 1 |
| 47 | chr1:22906049:T:C | NM_004442.7 | 10/16 | p.R611W | JM | 5.062E-05 | 1 |
| 48 | chr1:22906050:A:G | NM_004442.7 | 10/16 | p.R611Q | JM | 5.062E-05 | 1 |
| 49 | chr1:22906757:G:A | NM_004442.7 | 11/16 | p.R647G | TKD | 5.074E-05 | 1 |
| 50 | chr1:22906770:C:T | NM_004442.7 | 11/16 | p.V651A | TKD | 1.007E-04 | 2 |
| 51 | chr1:22906828:G:C | NM_004442.7 | 11/16 | p.S670R | TKD | 1.516E-04 | 3 |
| 52 | chr1:22906853:A:G | NM_004442.7 | 11/16 | p.D679N | TKD | 7.428E-03 | 148 |
| 53 | chr1:22906855:A:C | NM_004442.7 | 11/16 | p.D679E | TKD | 5.019E-05 | 1 |
| 54 | chr1:22906886:A:G | NM_004442.7 | 11/16 | p.V690M | TKD | 5.017E-05 | 1 |
| 55 | chr1:22906956:A:G | NM_004442.7 | 11/16 | p.R713Q | TKD | 1.556E-04 | 3 |
| 56 | chr1:22907975:C:T | NM_004442.7 | 12/16 | p.V721A | TKD | 4.987E-05 | 1 |
| 57 | chr1:22907986:T:G | NM_004442.7 | 12/16 | p.V725L | TKD | 4.987E-05 | 1 |
| 58 | chr1:22907998:T:C | NM_004442.7 | 12/16 | p.R729W | TKD | 4.974E-05 | 1 |
| 59 | chr1:22908007:A:G | NM_004442.7 | 12/16 | p.A732T | TKD | 2.487E-04 | 5 |
| 60 | chr1:22908076:A:G | NM_004442.7 | 12/16 | p.V755I | TKD | 1.988E-04 | 4 |
| 61 | chr1:22908091:A:G | NM_004442.7 | 12/16 | p.V760I | TKD | 4.970E-05 | 1 |
| 62 | chr1:22908136:A:G | NM_004442.7 | 12/16 | p.D775N | TKD | 4.979E-05 | 1 |
| 63 | chr1:22909043:C:T | NM_004442.7 | 13/16 | p.W793R | TKD | 5.126E-05 | 1 |
| 64 | chr1:22909117:A:G | NM_004442.7 | 13/16 | p.M817I | TKD | 4.990E-05 | 1 |
| 65 | chr1:22910409:T:C | NM_004442.7 | 14/16 | p.R845W | TKD | 4.964E-05 | 1 |
| 66 | chr1:22910452:A:T | NM_004442.7 | 14/16 | p.L859H | TKD | 4.979E-05 | 1 |
| 67 | chr1:22913532:G:C | NM_004442.7 | 16/16 | p.R976G | SAM | 1.522E-04 | 3 |
| 68 | chr1:22913670:A:G | NM_001309193.2 | 17/17 | p.A990T |  | 7.094E-04 | 14 |
| 69 | chr1:22913682:T:C | NM_001309193.2 | 17/17 | p.R994W |  | 1.013E-04 | 2 |
| 70 | chr1:22913689:T:C | NM_001309193.2 | 17/17 | p.T996M |  | 1.013E-04 | 2 |
| 71 | chr1:22913712:T:C | NM_001309193.2 | 17/17 | p.P1004S |  | 5.039E-05 | 1 |
| 72 | chr1:22913748:A:G | NM_001309193.2 | 17/17 | p.D1016N |  | 1.523E-04 | 3 |
| 73 | chr1:22913751:A:G | NM_001309193.2 | 17/17 | p.G1017R |  | 1.015E-04 | 2 |

**Supplementary Table 2:** ICD10 code and self-report code in UKB for neurological conditions.

| Disease | Self-report | ICD10 |
|---|---|---|
| Dementia or Alzheimer's disease | 1263 | F01, F02, F03, G30 |
| Parkinson's disease | 1262 | G20, G21 |
| Chronic degenerative neurological | 1258 | G23, G31, G32 |
| Guillain-Barré syndrome | 1256 | G610 |
| Multiple Sclerosis | 1261 | G35 |
| Other demyelinating disease | 1397 | G37 |
| Stroke or ischaemic stroke | 1081 | G463, G464, I64, I694 |
| Brain cancer | 1031 | C70, C71, D33 |
| Brain haemorrhage | 1491 | I60, I61, I62, I691, I692, I693 |
| Brain/intracranial abscess | 1245 | G060,G07 |
| Cerebral aneurysm | 1425 | I671, Q282, Q283 |
| Cerebral palsy | 1433 | G80, A521, A504, I64 |
| Encephalitis | 1246 | A83, A86, B011, B020, B262, A85, B004, B582, A84, B050, B941, G04, A321, G05 |
| Epilepsy | 1264 | G40, F803 |
| Head injury | 1266 | S07, T040 |
| Infections of the nervous system | 1244 | A80, A81, A82, A83, A84, A85, A86, A87, A88, A89 |
| Ischaemic stroke | 1583 | G45 |
| Meningeal cancer | 1031 | C70, C793 |
| Meningioma (benign) | 1659 | D33, D32 |
| Meningitis | 1247 | G03, A170, A171, A203, G01, G02, G00, G07 |
| Motor Neuron Disease (ALS) | 1259 | G122 |
| Neurological injury / trauma | 1240 |  |
| Spina bifida | 1524 | Q05, Q760 |
| Subdural haematoma | 1083 | P100 |
| Subarachnoid haemorrhage | 1086 | I60, S066, P103 |
| Transient ischaemic attack | 1082 | G45 |

**Supplementary Table 3:**
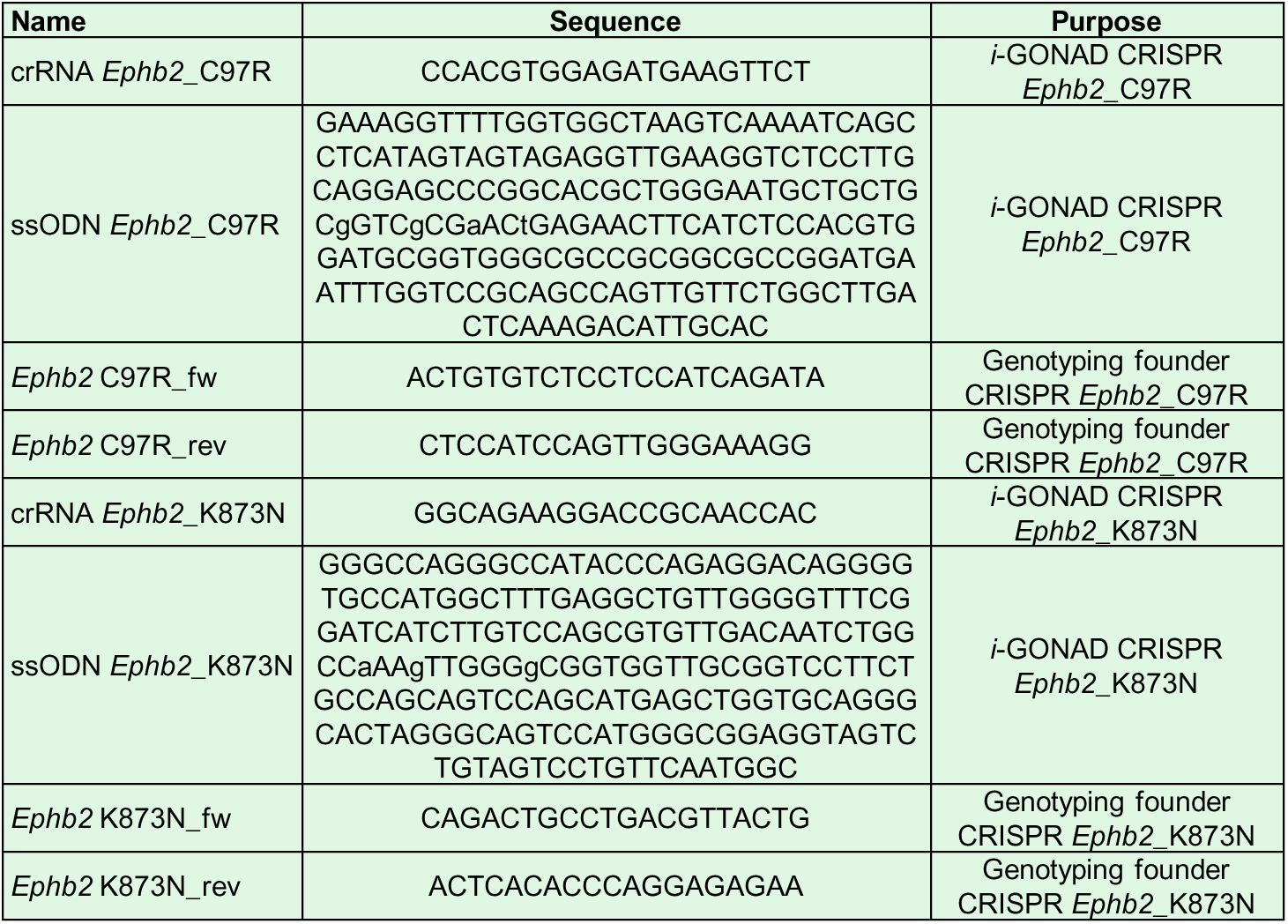
Primers used for generation and validation of EPHB2 knock-in mice.

